# Biophysical Basis of the *in vivo* Electroretinogram of the Mouse: 4-Shell Current Source Density Model

**DOI:** 10.1101/2025.01.28.635195

**Authors:** Gabriel Peinado Allina, Ratheesh Kumar, Vincent Yao, David J. Calkins, Paul FitzGerald, William E. Schiesser, Edward N. Pugh

**Affiliations:** Center for Neuroscience; Department of Ophthalmology & Vision Science; Department of Cell Biology & Human Anatomy; Department of Computational Biomedical Engineering, Lehigh University; Department of Physiology & Membrane Biology University of California, Davis; Department of Ophthalmology, Vanderbilt University

**Author notes:** Correspondence:* Edward N. Pugh, Jr., 3301 Tupper Hall, University of California, Davis, Davis, California 95616. Department of Neuroscience & Biomedical Engineering, Aalto University School of Science, Finland (Gabriel Peinado Allina).

## Abstract

A goal of contemporary physiology is to translate the knowledge obtained *ex vivo* of the molecular structure and function of ionic mechanisms into tools for quantitative, *in vivo* measurement of that function in humans, and in animal models of disease and therapeutic intervention. Non-invasive field potentials such as ECGs, EMGs, EEGs, and ERGs hold great promise in efforts to achieve this goal, but their full translation is challenging due to the multiplicity of cells with distinct ionic mechanisms and distributions of membrane current sources and sinks, and the requirement of adequate characterization of volume conduction in the relevant tissue(s). The molecular identities and subcellular distributions of the ionic mechanisms of mouse rod photoreceptors and the adjacent retinal pigment epithelium (RPE) have been thoroughly characterized, and are associated with two major components of the ERG, the *a*-wave and the *c*-wave, respectively. To develop a molecular-biophysical description of these components of the ERG, we have pharmacologically and genetically isolated rod photoreceptor-driven currents, created a 3D “4-shell” model of volume conduction in the mouse eye and extraocular tissue, and solved the Equation of Continuity for trans-photoreceptor layer and trans-RPE layer sources. Corneal and intraocular measurements of *a*- and *c*-waves are shown to reject the classic Rodieck-Ford electrical circuit model of the ERG, but found consistent with a 4-shell model having realistic values for extracellular conductivity in the eye and extraocular tissues. Our results and analysis explain the large variation across studies in maximal *a*-wave amplitudes and show how *a*-wave amplitudes exceeding 1 mV can be achieved.

## INTRODUCTION

The physiological function of electrically active cells *in vivo* is affected by their morphology, their ion channel expression patterns and the properties of the tissues in which they reside. Field potentials, including electrocardiograms (ECGs), electromyograms (EMGs), electroencephalograms (EEGs), the auditory brainstem response (ABR), electroolfactograms (EOGs), electroretinograms (ERGs) and others, provide a means of measuring electrical currents in their native milieu with minimal tissue disruption, and also afford the possibility of non-invasive observation of electrophysiological function *in vivo*. Field potentials are also invaluable for longitudinal assessment of physiological function in individual humans and animals, for example during disease progression and therapeutic intervention, as in widely employed clinical use of ECGs, EMGs, EEGs ABRs, EOGs ERGs. Consequently, field potentials that are interpretable in terms of the underlying currents of the tissues, cells and molecular mechanisms that generate them are of particular value in the translational era of biomedicine.

The ERG is a complex field potential triggered by light stimulation of the retina, and can be measured with a non-invasive corneal electrode. Since the discovery of the rod dark current by (Hagins et al., 1970), understanding of the generator currents underlying various components of the ERG has considerably advanced (Steinberg, 1985b; Pugh, 1998; Robson and Frishman, 1998, 2014). Recent advancements have come in particular from experiments on rodents, in which both genetic and pharmacological manipulations of molecular mechanisms are possible, and for which comparison between *ex vivo* transretinal ERGs and *in vivo* corneal ERGs is feasible (Green and Kapousta-Bruneau, 1999; Heikkinen et al., 2012; Vinberg et al., 2014; Turunen and Koskelainen, 2017).

A major outstanding issue in the application of corneal ERGs *in vivo* is the large variation between different studies in the reported amplitude of any specific component. Thus, for example, in a series of well regarded recent studies of normal mice, the maximum amplitude of the *a*-wave, generally accepted as originating in the suppression of the rod photoreceptor dark current, varied from 200 to 1000 μV (Supplementary Material, Table S1). Such variation is problematic for comparison of different studies, and worrisome inasmuch as it could arise from factors that directly alter neuronal function, such as indirect effects of anesthetic on intraocular pressure on vascular perfusion, or as direct action on photoreceptor ion channel function. A second related unresolved issue is whether the measured amplitudes of corneal ERG components can be quantitatively explained in terms of the retinal current generators and an “equivalent circuit” of the eye and extraocular tissues encompassing the reference electrode (Rodieck and Ford, 1969; Steinberg, 1985b), or whether a volume conduction model is required, and if so what tissue conductivities and boundary layer conductances are appropriate. A third outstanding issue is whether a single comprehensive analysis can account for the amplitude, kinetics and light-dependence of a second major ERG component that arises from photoreceptor activity, the *c*-wave, a trans-RPE potential originating in the decline in K^+^_o_ in the subretinal space during the rod and cone photoresponses (Steinberg et al., 1980; Steinberg, 1985a).

In this and two companion papers we use genetic and pharmacological tools to isolate rod and cone photoreceptor-generated field potentials (*a*-waves), and the trans-retinal pigment epithelium (*c*-wave) components of the corneal ERG in mice and apply volume conduction theory to the mouse eye and extraocular tissues. The resulting analysis, in addition to providing a foundation for absolute scaling between these two components of the ERG and their underlying generator currents, also explains why the recording methodology adopted in our investigations yields ERGs whose amplitudes are substantially larger than those of most previous studies, and provides a likely explanation of much of the variation in ERG amplitudes in other studies.

## METHODS

### Animals and experimental protocol

#### Mice and husbandry

All mice were cared for and handled with approval of the University of California, Davis, Institutional Animal Care and Use Committee and in accordance with National Institutes of Health Guidelines. Adult mice (2–4 mo) C57Bl6/J mice from Jackson Laboratories or in-house breeding were housed in a 12:12-h dark/light cycle (∼ 200 lux) and dark-adapted overnight for a minimum of 8 h before recording.

#### Anesthesia, mydriasis and proptosis

Mice were anesthetized under red light (>680 nm) with isoflurane (1.5%) and positioned on a regulated heating pad that maintained a core temperature of 37°C, which was monitored with a rectal probe. A mixture of phenylephrine and tropicamide (2:1) was applied to the corneal surface to achieve mydriasis, after which the eyelids were gently retracted to facilitate proptosis (Fig. 1). Conductive methylcellulose was applied to maintain corneal moisture during the recording and to ensure electrical contact with the corneal electrode. Care was used to restrict the spread of the gel to the surface immediately beneath and encircled by the recording electrode

**Figure 1.**
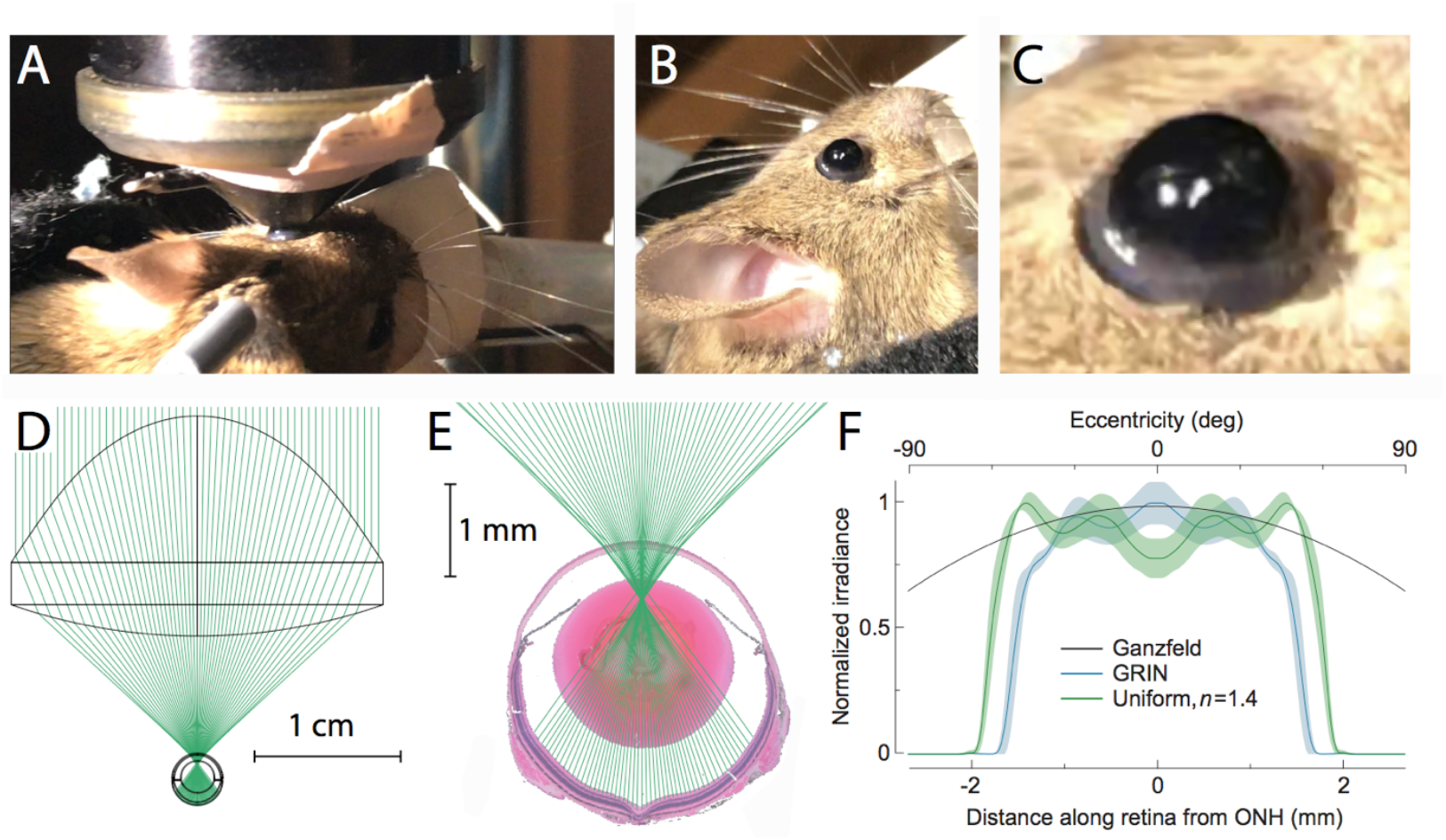
Characterization of the composite optical system formed by the Maxwellian view illumination and the mouse eye. **A**. Photo of anesthetized mouse with eye placed in position to receive light from the Maxwellian lens. **B**. Photo of the mouse in approximately the same position as in A but from a slightly different angle and with the Maxwellian lens removed. **C**. Close-up of the eye from panel B, showing the eye’s proptosis. The gray band somewhat below the equator of the ocular hemisphere is the conjunctiva, which is normally folded under the lid but revealed by the proptosis. **D**. Zemax^™^ raytrace model of the optical system comprising the Maxwellian lens and mouse eye’s optics, with rays from the LED filling the back aperture of the lens. **E**. Magnified display of the Zemax model showing the predicted paths of the rays, superimposed on an image of a section of a C57Bl6 mouse eye. **F**. Comparison of the power density of light at the retinal surface estimated for a Ganzfeld bowl illumination system (thin black line) or a Maxwellian illumination system. Zemax predictions are shown for models of the eye with a graded index of refraction (GRIN) model for the mouse lens (blue lines) and a homogeneous index of refraction (*n* =1.4; green lines). For the prediction of the retinal irradiance distribution from Ganzfeld illumination, a homogeneous refractive index (*n =* 1.4) was assumed for the mouse lens. The predicted irradiance distribution is spherically symmetric, and for a monochromatic source such as used ERG experiments would be scaled to the units of photons μm^-2^.

### Tonometry

The ERG affords the possibility of extracting information about the function of the neural and electrically responsive glial and epithelial tissues of the eye over epochs of an hour or more. A potential problem with the use of mydriatics is that they could alter the signal over time, for example, by elevating intraocular pressure, which can lead to “pressure blindness”. The effect of the pharmacological cocktail on intraocular pressure (IOP) was assessed by measuring the IOP of a cohort of mice every fifteen minutes during 1 hour with a Tonolab tonometer (Icare; Tampere, Finland), as previously described (Inman et al., 2006; Sappington et al., 2010).

### ERG apparatus

A modified commercial Maxwellian-view ERG apparatus (Phoenix Research Labs, Pleasanton, CA) was used for stimulation and recording. The corneal electrode of the Phoenix system was replaced with a custom-made 2 mm diameter platinum electrode mounted at the apex of a truncated cone 7.6 mm distance from the front surface of the Maxwellian lens (Fig. 1A, D). The electrode was shielded from the light stimulus by a thin black plastic cover to obviate photoelectric artifacts, and was aligned to the apex of the mouse eye under deep red (> 680 nm) light with the help of an infrared-sensitive CMOS camera aligned to the optical axis of the ERG apparatus (Fig. 1A). The edge of the dilated iris and the limbus of the eye were used as landmarks to guide the positioning of the corneal ring electrode.

### Zemax model of the eye and optics

The light stimulator included 508 nm (Osram #3397; Edmund Optics) and 365 nm Thorlabs, M365D1) LED sources, followed by a collecting lens positioned to deliver slightly convergent light that filled the back aperture of a Maxwellian-view objective lens (Edmund optics, #33-957) (Fig. 1D). In air (i.e., without the mouse emplaced) light from the sources was brought to a focal point in air 7.8 mm from the apex of the objective lens. To characterize the distribution of light on the retinal surface, we used the ray-tracing system Zemax™ to create a model of the complete optical system, including the mouse eye (Fig. 1, D, E). The Zemax model and software were used to estimate the power density (irradiance) distribution on the retinal surface and compare it with that of a Ganzfeld (Fig. 1F).

### Light calibration and stimulation protocols

Stimuli consisted of 1 ms flashes of light. The duration and the total energy delivered by the LEDs at the pupil plane or all stimuli were measured with a calibrated photodiode and calibrated neutral density filters. Flash strengths were expressed in units of photons µm^-2^ at the retinal surface, obtained by dividing total number of photons delivered per flash by a retinal surface of 1.8 ×10^7^ µm^2^ (18 mm^2^). As relatively intense flashes were used for this study, a minimum of 3 minutes in the dark was allowed for the retina to recovery between successive flashes.

### Electrical recording and post-processing

Recordings were made by a potentiometric differential amplifier provided with the Phoenix ERG system, with one input connected to the corneal electrode and the second to the reference electrode. The electrical system includes a 4-pole Bessel filter set to the widest bandwidth, corresponding to a high-pass cutoff frequency of 0.05 Hz and a low-pass cutoff of 1 Khz. Recordings were digitized at 5 KHz and saved on a hard disc. Data were post-processed with Igor Pro (WaveMetrics) scripts as indicated.

### Positioning and histological localization of the reference electrode

An important issue that must be resolved for a volume conduction model to predict the absolute amplitude of an ERG component is the locati on of the reference electrode in the extraocular tissue. Normally, the ∼ 250 µm diameter platinum needle that served as the reference electrode was inserted subcutaneously between bregma and the top of the pinna ipsilateral to the eye with the corneal electrode (Fig. 1A). The electrode was then gently advanced towards the eyeball until its tip was observed by small movements to be located near the superior eyelid. Care was taken to avoid rupturing the conjunctiva and entering into the peri-orbital fluid. To localize the reference electrode precisely in the extraocular tissue full-head histology was performed on two animals (Fig. 2). The mice were prepared for recording with the standard procedure described above. Following the application of phenylephrine and tropicamide and ocular proptosis, a 33 gauge insulin needle was inserted sub-epidermally in lieu of the reference electrode. The needle was used to inject a small quantity of China ink, labeling the path the electrode took as it is advanced towards the eye. Following the injection, the anesthetized animal was decapitated, and the head flash-frozen in isopentanol at -150° C. After 2 minutes, the head was transferred to an acidic acid (3%) and methanol (97%) solution at -60 C and subsequently stored at -80 C to allow the solution to penetrate the tissue. After one week, the head was mounted in a paraffin block and sliced into 200 µm thick sections. Sections were mounted in plastic, stained and imaged with a transmitted light microscope (Fig. 2). To illustrate the orientation of the 2D section with respect to the head, we used a 3D model of the mouse skull (Fig. 2A). The section with the electrode track corresponds to a plane roughly perpendicular to the bony orbit cavity’s widest dimension, displaced slightly coronally from the optic nerve head; and the closest proximity of the electrode track to the eye was ∼2 mm (Fig. 2B). (The distance of the electrode from the eye in the second experiment was approximately the same as in that illustrated.) In addition to its linear distance from the ocular surface, another defining characteristic of the reference electrode is its polar angle with respect to the corneal apex, described further below.

**Figure 2.**
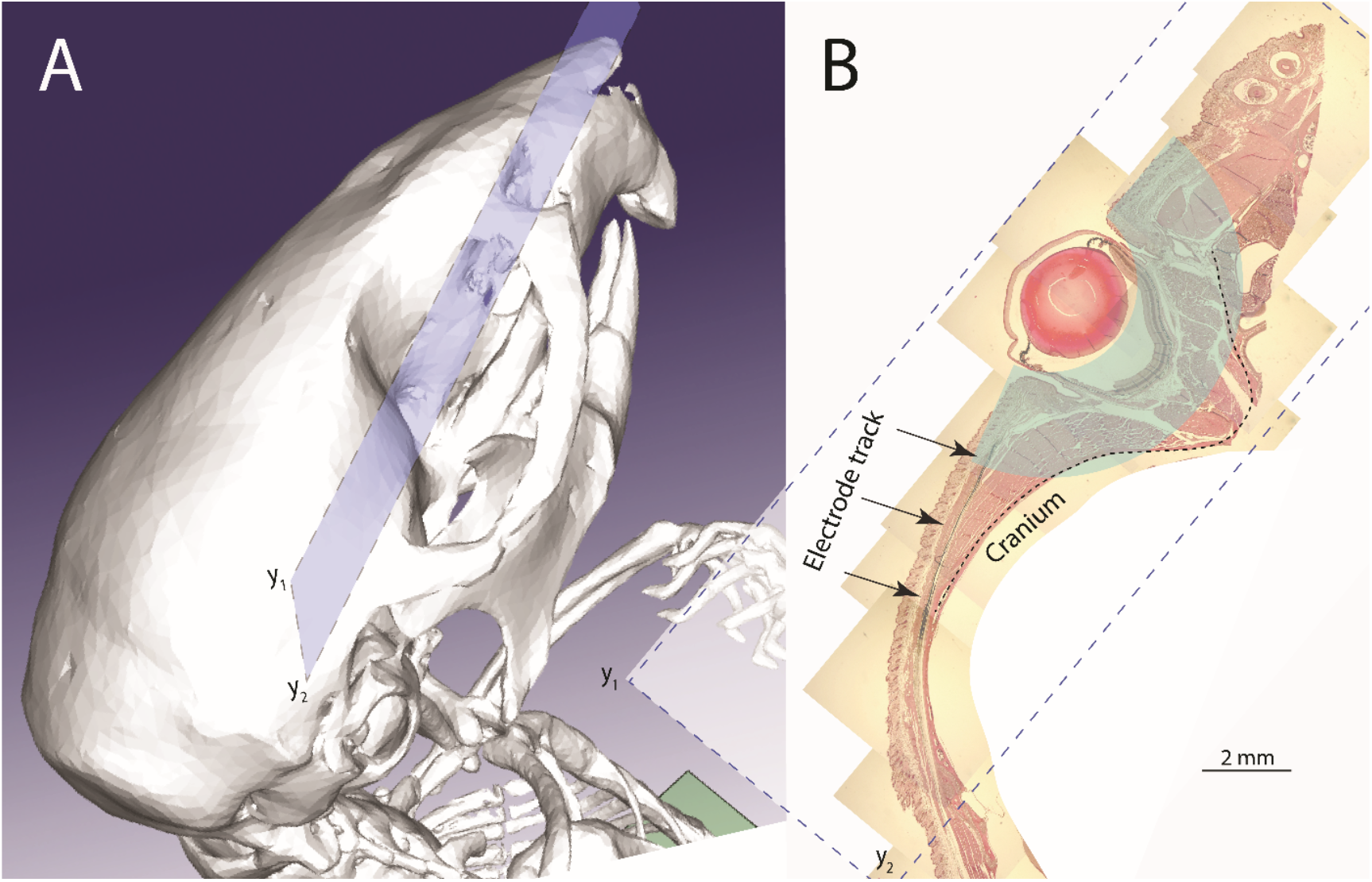
Structure of the skull and bony orbit of the mouse eye and histological localization of the reference electrode employed in ERG experiments. **A**. 3D rendering of the mouse skull arranged so that the viewer is looking down into the bony orbit in which the eye is situated. A blue parallelogram has been superimposed to indicate the approximate plane of section of two heads that were fixed for identifying the location of the reference electrode. **B**. Composite image of a section of a mouse head, with the plane of section illustrated in A. The subcutaneous track along which the reference electrode was inserted was marked with China ink. The pale green overlay highlights the region of extraocular tissue included in the volume current model; this region comes very close to the cranium, i.e., the bony orbit of the eye. In the fixation process the eye relapsed into the orbit; the green overlay identifies the extraocular region as it would be in the living, proptosed eye (Fig. 1C).

### Measurement of intraocular ERGs

In some experiments ERGs were measured intraocularly with a beveled, 200 µm diameter platinum wire serving as the electrode. The intraocular electrode was attached and aligned to the rotating shaft of a small (6 mm x 10 mm) motor. The motor speed was manually controlled by means of a potentiometer. The motor was mounted to a small micromanipulator and fastened to the ERG apparatus. The micromanipulator was aligned to the corneal electrode so that the recording electrode would penetrate the eyeball at the limbus (border between cornea and sclera). The motor was used to bore a hole, rather than simply puncture the eye, avoiding sharp changes in elastic tension, unnecessary eye movement and preserving alignment. When the intraocular electrode was employed, a switch was used that enabled alternating recording of corneal and intraocular ERGs in response to identical light stimuli with the same reference electrode. Intraocular ERGs were digitized and recorded in the same way as corneal ERGs.

## VOLUME CONDUCTION THEORY AND 4-SHELL MODEL OF THE MOUSE EYE

### Problems with the classic Electric Circuit Model of the corneal ERG and rationale for a volume conduction model

It is instructive to compare current flow in the classic Electrical Circuit Model (Fig. 3B) of the ERG (Rodieck and Ford, 1969; Steinberg, 1985b) with that expected in a volume conduction model of the eye (Fig. 3C). The Circuit Model is topologically equivalent to flattening the hemispherical ocular tissues and embedding them as a layer cake in a cylinder with a non-conductive wall (Fig 3A). Extracellular current flowing across any specific retinal layer generates a corresponding translayer potential, which is incorporated into the model as an ideal voltage source connected in parallel to a resistor representing the trans-layer extracellular resistance, and in series to the rest of the circuit. Thus, for example, the extracellular limb of the rod dark current causes the photoreceptor synaptic layer (i.e., the outer plexiform layer or OPL) to have a positive potential with respect to the photoreceptor outer segment tips (Hagins et al., 1970). This translayer potential drives current across the photoreceptor layer of the retina, and also around the full loop of the eye ball causing the cornea to be at a positive potential relative to a reference electrode in the extraocular return path, and giving rise to the corneal-negative *a*-wave when the dark current is suppressed by light exposure.

**Figure 3.**
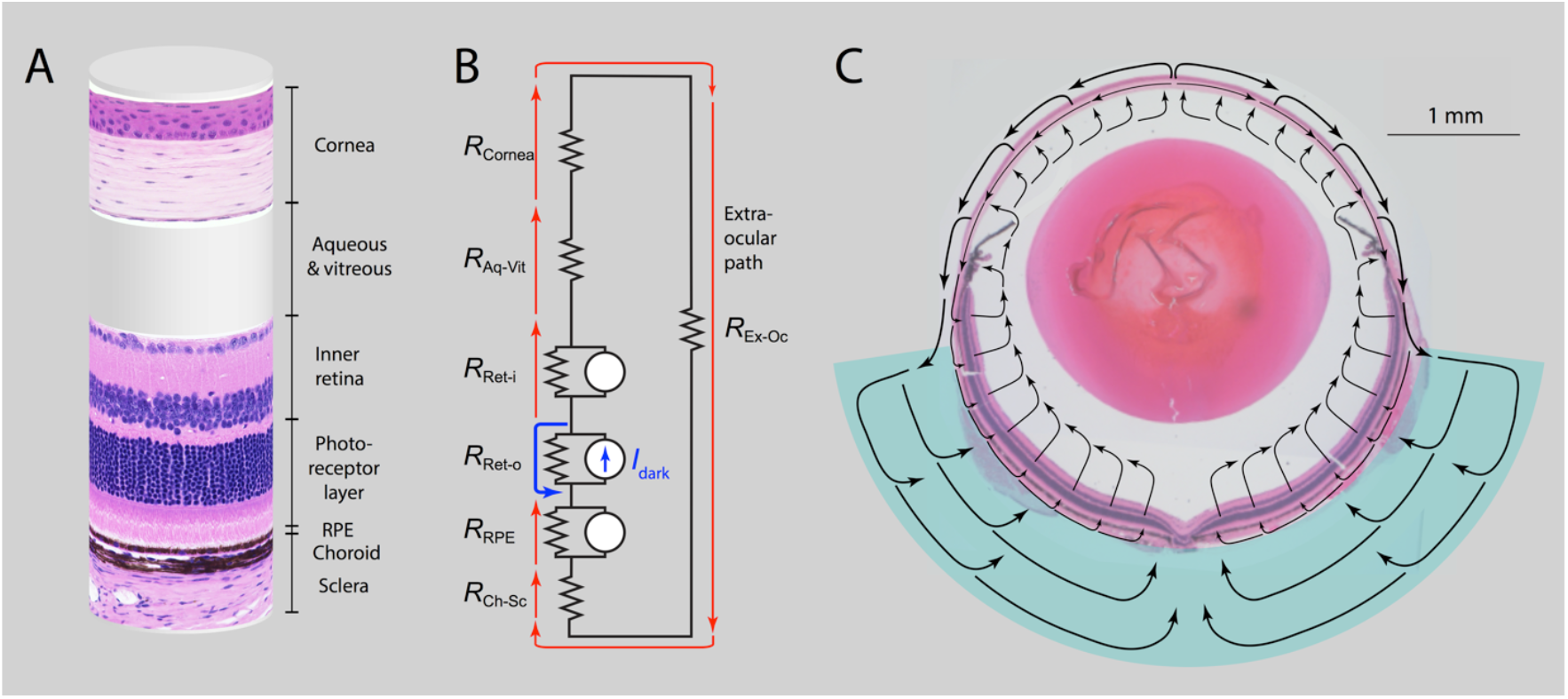
Comparison of current flow in the classic electrical circuit model of the ERG with that in a volume conduction model of the mouse eye. **A, B**. Rodieck-Ford electrical circuit model of the ERG. The electric circuit model effectively topologically distorts the retinal tissues that generate field potentials and the auxiliary tissues and media through which intraocular current flows into an insulated, radially uniform cylinder. Different ocular layers have different resistances identified in B with “*R*_X_” where X is an abbreviation for the layer or structure identified in the cylindrical structure. In B, radially extended neurons and glial cells in different layers generate axial currents that flow in a loop within the layer, with current flowing axially interior to the cell in one direction, and exteriorly in the opposite direction: these translayer currents give rise to translayer potentials that in turn drive current around the entire loop, which includes a poorly specified extraocular path. The dark current *I*_dark_ of the photoreceptor layer (PRL) is singled out as a source, and the current flow in the overall loop corresponds to that caused by the trans-PRL potential created by *I*_dark_. In the model, the ERG is measured across the extra-ocular resistance or some part thereof. **C**. Schematic of volume current flow in the mouse eye and extraocular tissue expected from a trans-photoreceptor layer source imposed on an image of the mouse eye. The eye is presented as it would be in the experimental conditions of the ERG recordings, i.e., proptosed relative to the head and extraocular tissue (cf. Fig. 1); the region of the extraocular tissue is illustrated in panel C in pale green, but drawn to scale (cf. Fig. 2B).

Treating the eye as a multi-layer, spherical structure through which currents generated by retinal layer sources flow (Fig. 3C) makes it possible to address important issues that are eschewed by the Circuit Model. For example, how does the spatial inhomogeneity of the retina -- as manifest in the systematic variation of ONL thickness and thus photoreceptor density – affect the intraocular potential in the ocular media? Are the intraocular media isopotential, and if not does the intraocular potential vary in a manner that affects the amplitude of ERG components? Is the posterior pole (RPE, choroid, sclera) of the eye isopotential with respect to polar angle, and if not how does this affect the return current paths and the amplitudes of ERG components? Where should the reference electrode be placed so as to achieve maximum amplitude of ERG components?

To address these and other issues we developed a biophysically based volume current model of the mouse eye and extraocular orbital tissue (Fig. 4) and applied it to measurements of two principal ERG components, the *a*-wave and the *c-*wave. The complete model is presented in three papers. This first paper characterizes the volume currents and their corresponding potentials in the non-retinal ocular tissues, including the cornea, sclera, conjunctiva and post-orbital tissue generated by a trans-photoreceptor retinal layer (*a*-wave) or a trans-retinal pigment epithelium (RPE; *c*-wave) source. The second paper predicts the absolute magnitude, axial profiles and kinetics of the extracellular currents of rod photoreceptors and their transretinal potentials, and with the 4-shell model predicts the corresponding time- and flash-strength-dependent corneal *a*-waves. The third paper predicts the absolute magnitude and kinetics of the *c*-wave, based on the hypothesis that it arises from extracellular changes in the concentration of K^+^ in the subretinal space consequent to the photoreceptor light response acting on Kir channels in the RPE apical membrane.

**Figure 4.**
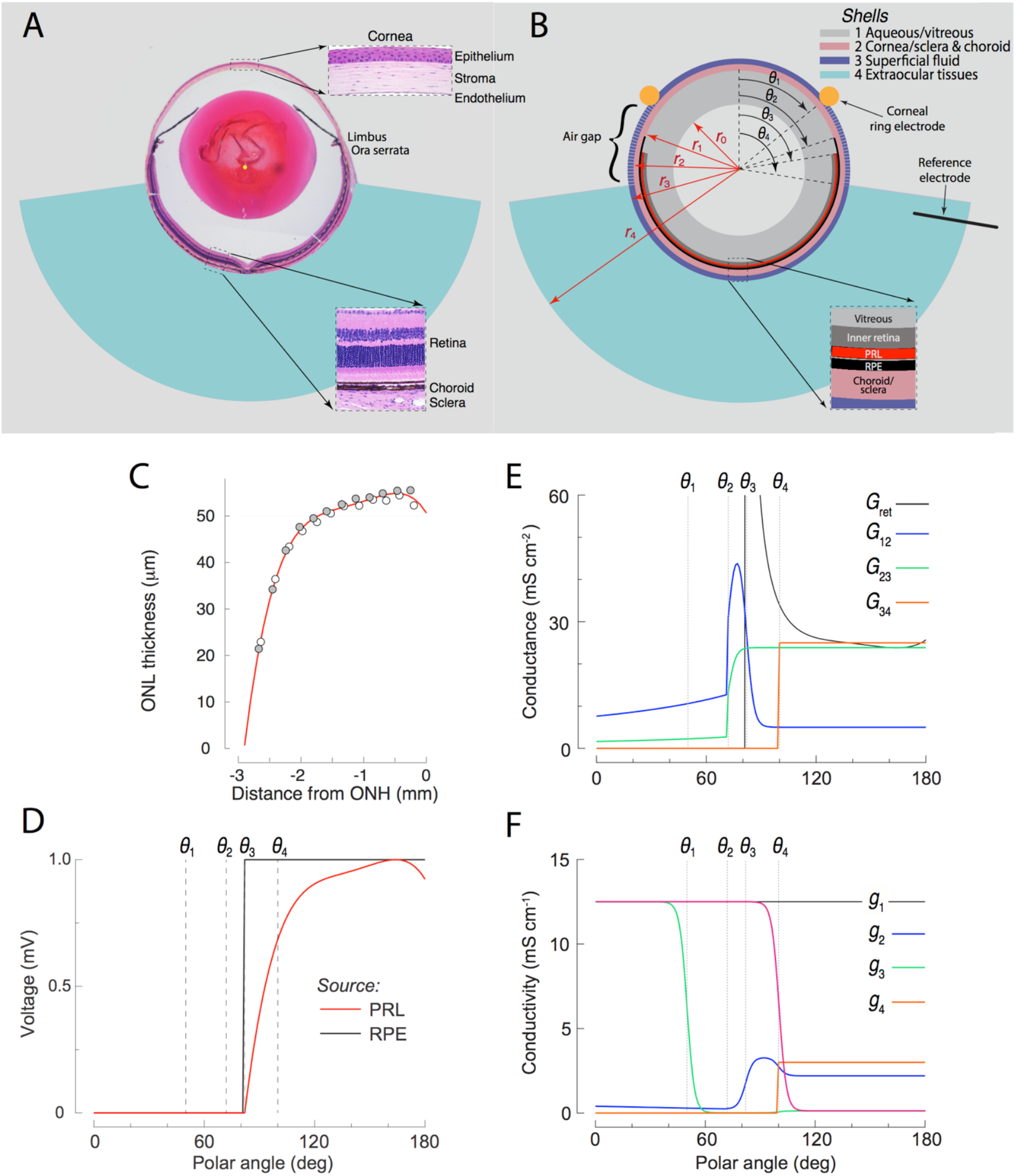
Distributions of the layer conductivities, boundary conductances and photoreceptor layer (PRL) and retinal pigment epithelium (RPE) sources in the 4-shell model of the mouse eye. **A**. Section of a mouse eye positioned with respect to the schematic of the extraocular tissues, drawn to scale. The insets provide magnified histological images of the cornea, and of the retina-RPE-choroid-sclera layers of the posterior eye. **B**. Schematic of the spherical polar coordinate system with identification of key dimensions, and radial and angular transitions. The lens is treated as spherical with radius *r*_0_ and is centered in the coordinate system. In order outward from the lens the shells are as follows: the aqueous and vitreous humors (Shell 1; thickness *r*_1_ − *r*_0_); the cornea (anteriorly) and the choroid and sclera (posteriorly) (Shell 2; thickness *r*_2_ − *r*_1_); the superficial fluids bathing the eye (Shell 3; thickness *r*_3_ − *r*_2_); the extraocular tissues (Shell 4; thickness *r*_4_ − *r*_3_). The apex of the cornea forms the north pole of the coordinate system. Angles at which key transitions of media or tissue type occur are identified: *θ*_1_ (50 deg) angular position of corneal ring electrode; *θ*_2_ (72 deg), end of cornea and beginning of ora serrata; *θ*_3_ (83 deg), end of ora serrata and angular beginning of retina; *θ*_4_ (100 deg), angular beginning of extraocular tissues. **C**. Thickness of the outer nuclear layer as a function of the distance along the retinal surface from the optic nerve head (ONH). The data points (symbols) are taken from (Wu et al., 2014): the original data extended in both directions from the ONH, and those extending to the left were reflected about the ONH; the data are well described by 4^th^-order polynomial (red curve). **D**. Angular distributions of the photoreceptor layer (PRL, red curve) and retinal pigment epithelium (RPE, black line) sources in the model. The PRL source distribution is identical to the red curve in panel C, except replotted with respect to the polar angle coordinate system of the model. The RPE source distribution is a step function, based on the assumption that the RPE layer properties are invariant across the retina. The ONH forms the south pole (*θ* = 180 deg) of the polar coordinate. **E**. Polar angle distributions of the conductances *G*_j,j+1_(*θ*), j=1,3 of the boundaries between the shells (cf. Eqs 3.1-3.4 and Eq 4). **F**. Polar angle distributions *g*_k_(*θ*) of the conductivity within each shell (*k* = 1-4).

### General principles of volume conduction in living tissues: current source density (CSD) analysis

#### Down the gradient

The extracellular spaces of living tissues are volume conductors in which natural magnetic fields are negligible. Accordingly, electrical currents in tissues obey two fundamental relations, the first of which is expressed as follows:

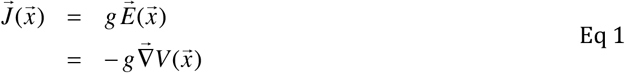

Here 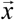 is position, 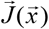 current density (units: A cm^-2^), *g* conductivity (units: S cm^-1^), 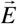 the electric field (units: V cm^-1^) and *V* the electric potential (V, volts). Equation 1 expresses the idea that current density is a position-dependent vector quantity whose magnitude is proportional to and direction is the same as that of the electric field. The conductivity *g* is generally position-dependent and, if the tissue is anisotropic, can be a tensor (3 × 3 matrix). The second line of Eq 1 embodies the insight that in a such a volume conductor there is a unique position-dependent scalar potential 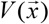 whose negative gradient defines the electric field: the negative sign embodies the insight that current flows “down the gradient,” or equivalently, that path-independent work is performed in moving a charge from a lower to a higher level of the potential in the volume.

#### Continuity

The second general relation obeyed by extracellular currents in a volume conductor is

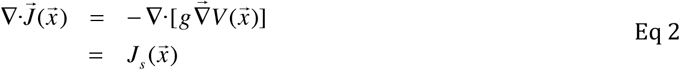

Equation 2 is a variant of the Equation of Continuity (EOC), and is sometimes referred to as Charge Conservation in the context of volume current theory. In the left hand side (LHS) of the first line of Eq 2 the divergence operation 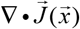 computes the sum of the derivatives of the components of current flowing into and out of the volume element in the three orthogonal directions of the coordinate system, while the first line of the right hand side (RHS) sub-stitutes the expression for 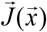 from Eq 1. The volume current density 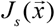 (units: A cm^-3^) is a scalar quantity, and in the elementary volume (voxel) at location 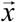 must be zero unless there is a source that delivers current into the voxel, or a sink that extracts current. In living tissues, current sources and sinks are created by charges flowing across (ionic) or onto (capacitive) membranes of electrically active cells. In our analysis we will also use the formalism of sources and sinks to describe currents that flow between distinct adjacent tissues. The application of Eq 2 to deduce membrane currents from field potentials is known as current source density (CSD) analysis (Nicholson and Freeman, 1975; Buzsaki et al., 2012; Einevoll et al., 2013). A classic application of CSD analysis in a rat retina slice preparation led to the discovery and initial characterization of the rod photoreceptor dark current (Hagins et al., 1970).

### Features of a 4-shell volume conduction model of the mouse eye

The model treats the mouse eye as comprising four concentric spherical shells with varying conductivities and boundary layers surrounding an effectively non-conducting spherical inner core, the crystalline lens (Fig. 4B). The overall goal is to predict the potential distribution 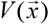 (Eq 2) throughout the ocular and periocular tissues arising from a specified trans-photoreceptor layer or retinal pigment epithelium (RPE) source. Specifically, we sought to derive the potential difference between the corneal and reference electrodes, with the location of the reference electrode in the extraocular tissue experimentally determined, and with variations in the way the corneal electrode is situated on the eye. The rationale for a concentric spherical shell model is that the eye itself is spherical, and its tissues exhibit relatively abrupt changes in conductivity at specific radial and angular distances from the ocular center allowing these transition to be treated as boundary layer conductances. (For the most part, we adopt CGS units for their familiarity and applicable scale, but always report the units.)

#### Polar coordinate boundaries of the mouse eye model

The volume conduction model employs a spherical coordinate system with the apex of the cornea as the north pole (*θ* = 0) and the optic nerve head as the south pole (*θ* = 180 deg) (Fig. 4B). The model assumes azimuthal homogeneity. The model specifies the key polar angles at which various transitions occur, including the angular extent of the spherical cap of the cornea encompassed by the ring electrode (0 ≤ *θ* ≤ *θ*_1_), the angular extent of the cornea itself (0 ≤ *θ* ≤ *θ*_2_), the ora serrata (*θ*_2_ ≤ *θ* ≤ *θ*_3_), and the extent of exposed conjunctiva in the proptosed eye (*θ*_2_ ≤ *θ* ≤ *θ*_4_) . Conductivity varies amongst the different component tissues and as a function of polar angle, as do the conductances of the boundary layers separating the shells. Hence, before presenting the mathematical description of model, we first describe the relevant features of the eye and extraocular tissues (Table 1).

**Table 1:**
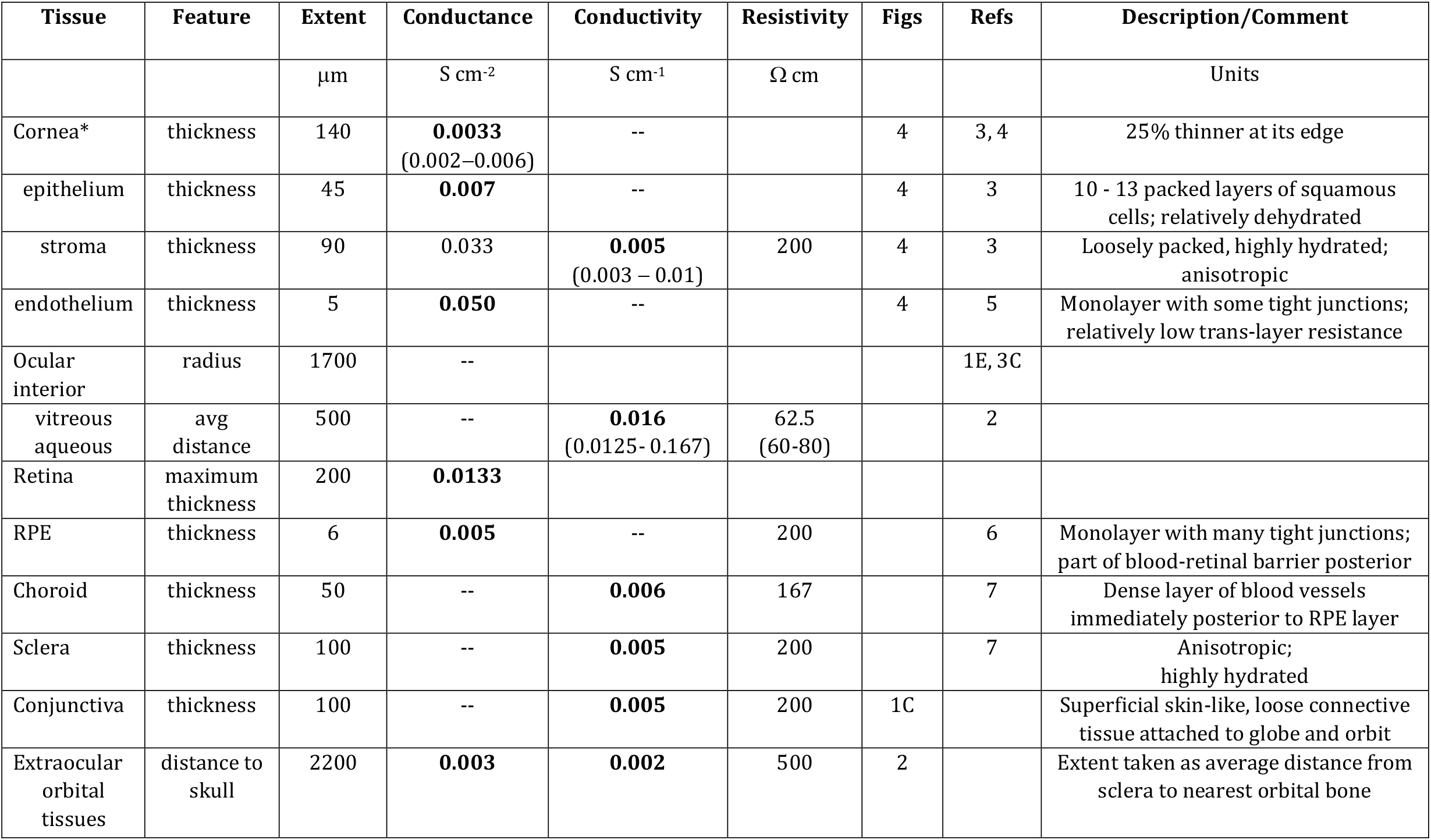
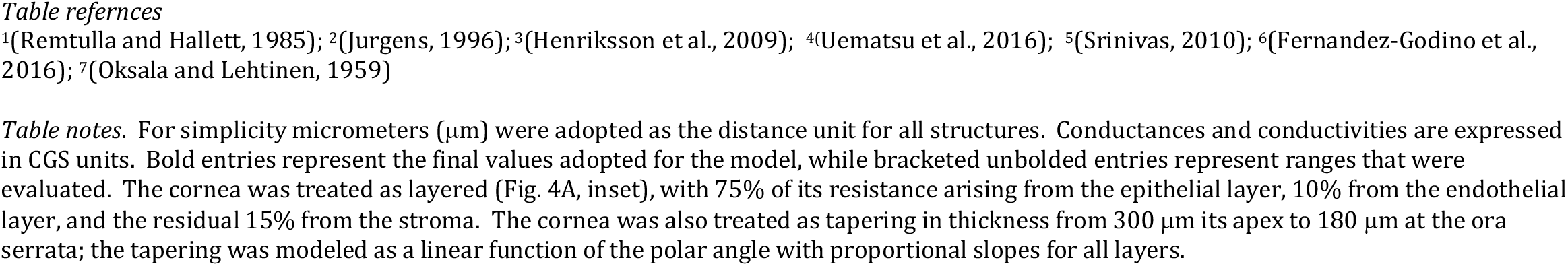
Dimensions and Electrical Properties of Mouse Ocular and Extraocular Tissues.

#### Photoreceptor and RPE retinal layer sources; retinal layer resistances

This paper develops and tests predictions of a volume current model for field potentials in the eye and extra-ocular tissues arising from the trans-photoreceptor layer potential (*V*_PRL_) arising from the dark current, or from trans-epithelial potentials of the RPE (*V*_RPE_.) In contrast to the implicit assumption of the conventional circuit model that the retina is spatially homogeneous (Fig. 3A, B), *V*_PRL_ is treated here as depending on retinal eccentricity (Fig. 4D), due to variation rod density, as manifest in the thickness of the layer of photoreceptor nuclei, the ONL (Fig. 4C). In both cases, the source potentials are assigned the nominal value 1 mV. In the companion papers the absolute magnitudes of the source potentials will be derived from current source density analysis of the respective layers. Compared with the vitreous humor (Shell 1), the retina has negligible thickness (< 200 μm). However, the resistance of the fraction of the retina lying vitreal to the PRL or RPE sources is material. We estimated the overall resistance of the ∼200 μm thick mouse central retina to be 75 Ω cm^2^, and assumed that the resistance as a function of polar angle to be inversely proportional to thickness. The corresponding angular conductance distribution, *G*_ret_ (*θ*), is plotted in Fig. 4E for the PRL source. When the source is the RPE, the substantial resistance of the PRL (42 Ω cm^2^ in the central retina) was included in the distribution.

#### Crystalline lens

The crystalline lens of the mouse fills much of the ocular interior, and is roughly spherical with its center displaced forward from the center of the eye (Fig. 4A). In the model, the lens is treated as a non-conducting sphere of radius *r*_0_ = 0.11 cm centered in the eye (central light gray sphere in Fig. 4B).

#### Shell 1: Intraocular media

The aqueous and vitreous humors comprise a medium of high conductivity between the lens and the interior tissues of the ocular globe (Table 1). The resting (dark adapted) vitreal-positive trans-photoreceptor layer potential *V*_PRL_ drives current into the vitreous (Fig. 3C): some of this current flows into the cornea (0 ≤ *θ* ≤ *θ*_2_) and into the portion of the sclera that does not have underlying retina and RPE (*θ*_2_ ≤ *θ* ≤ *θ*_3_) . The current will exit the eye via electrolyte bathing the cornea, and through the interior of the corneal layers, trabecular meshwork, and ora serrata and sink across the sclera, choroid, and RPE to the outer segment side of the photoreceptor layer, or to a general ground in the head and body (Fig. 4A).

#### Shell 2: Cornea, sclera and choroid

As revealed by the analysis, volume conduction interior to the multilayered cornea is a critical feature that must be captured in a model of the ERG. The mouse cornea (Fig. 4A, upper inset) is ∼ 140 μm thick at its apex, thinning to about 100 μm at its extreme (Henriksson et al., 2009). The cornea defines the north polar region of Shell 2 (0 ≤ *θ* ≤ *θ*_2_), and comprises three distinct layers. A thin endothelium forms the innermost layer, a highly hydrated stroma with collagen fibers the middle layer, and a layered epithelium the outermost layer. Histology reveals a major difference between the relatively dehydrated squamous cell outer layer of the epithelium, and the cuboidal cells of the inner layer, manifesting substantially different layer conductances (Table 1). A positive electrical potential in the aqueous drives current through the relatively conductive corneal endothelium into the stroma (Fig. 4A). Current entering the cornea can flow “downhill” toward the sclera through the corneal stroma, or outward across the epithelium to the corneal surface, whence it can then flow into extraocular tissues if the cornea is covered by tears or gel. The combined sclera, choroid and conjunctiva (*θ*_2_ ≤ *θ* ≤ *θ*_4_), or simply sclera and choroid (*θ* ≥ *θ*_4_) comprise the remainder the shell. Shell 2 is treated as a “thin” layer, meaning that the potential is assumed to have no radial variation across the layer. As mentioned above, the model also assumes there to be no azimuthal variation in the shell at any given polar angle *θ*.

#### Shell 3: Superficial fluid layer

This shell comprises a layer of conductive gel or tears on the corneal surface, and a layer of secreted extracellular fluid behind the eye, posterior to the closed conjunctival sac (Fig. 4B, dark blue band). Our standard recording protocol introduces an air gap outside the corneal ring electrode (Fig. 4B), a manipulation expected to substantially increase resistance in Shell 3 between polar angles *θ*_1_ and *θ*_4_, thereby decreasing the current flow in this shell. In some experiments we intentionally introduced highly conducting gel in this polar region to test the hypothesis that the air-gap is a key factor underlying the high amplitude of the ERGs we record. Shell 3, like Shell 2, is treated as a thin layer, and is thus assumed to have no radial variation in potential, and likewise to have no azimuthal variation.

#### Shell 4: Extraocular tissues

In the proptosed eye (Fig. 1C), Shell 4 is approximately 2.2 mm thick, and spans all the extraocular tissues up to the bony orbit of the skull (Fig. 2B and Fig. 4B, bluish-green region). Shell 4 comprises a heterogeneous group of tissues that includes skin-like connective tissue, the extraocular muscles, the optic nerve and vasculature. An essential feature of Shell 4 is that it extends far enough from the eye to encompass the reference electrode ring (Fig. 2B; Fig. 4B). A second feature is that it (unlike Shells 2 and 3) includes a radial coordinate grid. This radial grid enables the model to quantify the potential variation in the extraocular tissues at distances up to and beyond the position of the reference electrode.

Two limitations of Shell 4 as a physical model of the eye are that the conductivity of the extraocular tissues is assumed constant (Fig. 4B; Table 2), and that the shell is treated as radially bounded, whereas the extraocular tissues comprise heterogeneous conductive paths through and around the skull (Fig. 2). The first of these assumptions is unavoidable because the variation in tissue type is on a fine scale that cannot be readily determined in 3D with submillimeter resolution. The effect of this assumption can be to some extent be tested by varying the conductivity over a range that includes measured values for the different tissue types. The second limitation is mitigated by results predicting that the potential decays rapidly with radial distance from the posterior surface of the eye and is essentially negligible at the outer boundary of the shell.

**Table 2:** Structural Properties of the 4-Shell Volume Conductor Model of the Mouse Eye and Extraocular Tissues.

| Description | Symbol | Units | Values | Equations | Figure |
| --- | --- | --- | --- | --- | --- |
| Radius of the crystalline lens | $r_0$ | cm | 0.11 | 3 | Figs. 4A, B |
| **Shell 1:** Vitreous/aqueous between lens surface and corneal endothelium or retinal surface | $\Delta r_1$<br>($r_1 - r_0$) | cm | 0.06 | 3.1 | Fig. 4B |
| **Shell 2:** Cornea (anterior 71 deg)<br>sclera+choroid (posterior 109 deg) | $\Delta r_2$<br>($r_2 - r_1$) | cm | 0.011 | 3.2 | Fig. 4B |
| **Shell 3:** Conducting fluid layer<br>on ocular surface<br>subconjunctival | $\Delta r_3$<br>($r_3 - r_2$) | cm | 0.030<br>0.010 | 3.3 | Fig. 4B |
| **Shell 4:** Region behind eye to bony orbit | $\Delta r_4$<br>($r_4 - r_3$) | cm | 0.22 | 3.4 | Figs. 2, 4B |
| Conductivity of Shell j | $g_j(\theta)$ | S cm<sup>-1</sup> | Polar angle-dependent | 3,16 | Fig. 4E |
| Conductance of boundary between shells j and j+1 (j = 0, ..., 3) | $G_{j,j+1}(\theta)$ | S cm<sup>-2</sup> | Polar angle-dependent | 3 | Fig. 4F |

*Table notes.* The cornea was treated as tapering from its apex to its edge (see notes to Table 1).

### Mathematical Realization of the 4-Shell Model

As the eye and its concentric layers are approximately spherical, Eq 2 was expressed in spherical coordinates, resulting in the following partial differential equation (PDE) for the potential *V* as a function of polar angle *θ* and the radial distance *r* from the center of the eye:

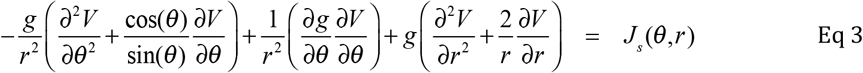

Equation 3 is the direct transformation of Eq 2 into polar coordinates subject to the constraint of no azimuthal variation, i.e. that *V* exhibits only polar and radial variation.

The division of the model into concentric shells with varying conductivities and inter-shell boundary conductances makes it useful to express Eq 3 separately for each shell. This approach allows the introduction of explicit expressions for source/sink terms at tissue boundaries or transitions. For convenience *V*_i_ (*i* = 1 − 4) is used to represent the potential in Shells 1 to 4, respectively, with *V*_1_, *V*_2_ and *V*_3_, being functions only of the polar angle *θ*, while *V*_3_ is a function of both *θ* and *r*. The specific form of Eq 3 applicable to each of the shells is now presented, followed by a brief description.

#### Shell 1 (aqueous; vitreous)

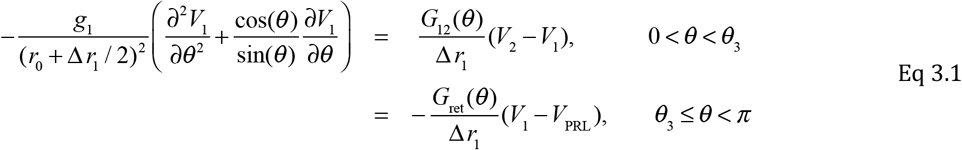

In Eq 3.1 *r*_0_ is the radius of the lens (Fig. 4B), while Δ*r*_1_ = *r*_1_ − *r*_0_ is the thickness of the shell, and *g*_1_ its conductivity, assumed homogeneous. For *θ* < *θ*_3_ the right-hand side (RHS) of Eq 3.1 describes volume current flowing between the aqueous and the cornea, trabecular meshwork and ora serrata: while both *V*_0_ and *V*_1_ are functions of *θ*, generally *V*_2_ < *V*_1_ in this domain of the polar coordinate. For *θ* ≥ *θ*_3_ the RHS of Eq 3.1 describes volume current entering Shell 1 driven by the retinal source *V*_PRL,_ taken to be the vitreal-positive potential generated by the dark current of the photoreceptor layer (PRL). In Eq 3.1 *G*_ret_ is the polar angle-dependent conductance of the retina anterior to the photoreceptor layer, while *G*_12_ is the polar angle-dependent conductance (units: S cm^-2^) between layers 1 and 2 (Fig. 4E) and the product *G*_12_ (*θ*)[*V*_2_ (*θ*) − *V*_1_ (*θ*)] specifies the radial current density (A cm^-2^) across the boundary between Shells 2 and 1 at the specific polar angle. Division of the RHS by Δ*r*_1_ is required to convert the source term into a volume current density (A cm^-3^), as explained in the subsequent section “Tissue conductivities and boundary layer conductances.” An important feature of Eq 3.1 is that it explicitly incorporates retinal eccentricity-(polar angle-) dependence of the retinal source, as *V*_PRL_ is specified as a function of *θ* (Fig. 4C, D). (The boundary conditions at the poles (*θ* = 0, *θ* = *π*) for all four shells are presented together below.)

#### Shell 2 (cornea; sclera+choroid)

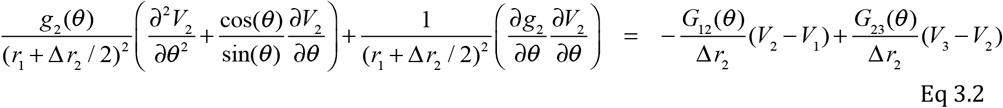

In Eq 3.2 *r*_1_ is the radial distance from the center of the eye to the inner boundary of Shell 2, which in the north polar region is the corneal endothelium and in the posterior pole the RPE, each of which is less than 10 μm thick. The radial distance to the center of Shell 2 is *r*_1_ plus half the thickness of the shell, i.e., *r*_1_ + Δ *r*_1_ / 2 . The conductivity *g*_2_ (*θ*) of Shell 2 varies as a function of polar angle (Fig. 4F), a variation that affects both terms on the LHS of Eq 3.2. The RHS of the equation represents source/sink terms at the inner and outer boundary of the shell, respectively. In the polar region without retina *θ* < *θ*_3_ the first term on the RHS represents current flowing between the cornea and the vitreous, while the second term describes current flowing between the corneal epithelium and the superficial bathing fluid. Because *V*_1_ > *V*_2_ and *V*_2_ > *V*_3_, these two terms describe a source and sink, respectively. In the polar region where there is retina *θ* ≥ *θ*_3_, the conductance boundary between Shells 1 and 2 is formed by the RPE, and the potential on the retinal side of the RPE is zero for the photo-receptor layer dark current source: in this case, the first term on the RHS is simplified and acts as a sink for Shell 1, while the second term would also be expected to describe a sink. As in Eq. 3.0, division by the shell thickness Δ *r*_2_ is required to correctly convert the source/sink terms into volume current density (A cm^-3^).

#### *Shell* 3 (*corneal tear/gel layer; post-ocular fluid layer*)

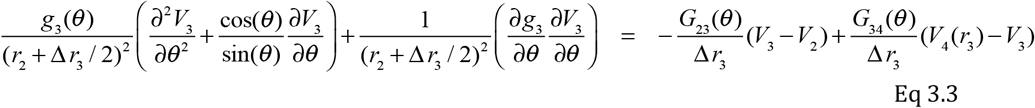

In Eq 3.3 *r*_2_ is the radial distance to the inner boundary of Shell 3, and as for Shell 2, the radial position is taken as distance to the middle of the shell: *r*_2_ + Δ *r*_3_ / 2 . The terms on of Eq 3.2 are parallel to those in Eq 3.1, including both a source at the inner boundary, and a sink at the outer boundary.

#### Shell 4. Extraocular tissues

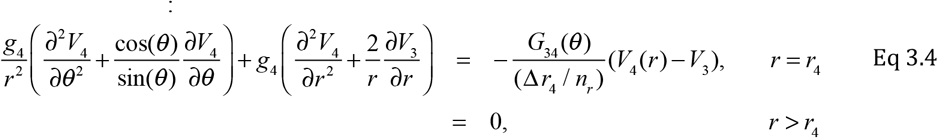

In Eq3.4 *r*_3_ is the radial distance to the inner boundary of Shell 4, but *r* varies in this shell over the range *r*_3_ ≤ *r* ≤ *r*_4_ (bluish-green region in Figs. 2B, 3C, 4A, 4B), necessitating inclusion of divergence terms with first and second derivatives with respect to *r*. There is a source/sink term at the inner boundary of Shell 3 (*r* = *r*_*3*_), but otherwise the volume current density is source/sink-free. Shell 4 has thickness Δ*r*_4_ = *r*_4_ – *r*_3_ and is radially subdivided into *n*_*r*_ sub-shells, whose individual thickness is Δ*r*_4_/*n*_*r*_. Source current flowing into Shell 4 moves down both polar and radial gradients. The RHS of Eq 3.4 treats each subshell as radially isopotential, and flow between the subshells follows the gradient. The model is assumed closed, so that no current flows beyond Shell 3.

### Tissue conductivities and boundary layer conductances

Ideally, a biophysical model involves no free parameters, i.e., employs only dimensions and parameter values known from prior research or obtainable from measurements. While the dimensions of the mouse eye and extraocular tissues are known (Figs. 1-4), the values of its tissue conductivities and layer conductances have not been as thoroughly investigated, and so call for additional explanation and justification. *Conductivity* is an intrinsic, local property of a tissue or medium: it is the proportionality factor between local electric field strength and current density (Eq 1) at a specific spatial location; from unit analysis conductivity is readily seen to have the CGS units S cm^-1^. *Conductance* is a property of a spatially extended layer of material or tissue such as the cornea and RPE and has CGS units S cm^-2^. (Usually, when measurements are made of isolated tissue sheets, they are reported as *resistance* (CGS units: Ω cm^2^), reciprocal to conductance.) If a conducting layer of thickness Δ*L* has a uniform structure and conductivity *g*, the conductance *G* of the layer is related to the conductivity by *G* = *g /*Δ*L*.

The boundary layer conductances *G*_ij_ in Eqs. 3.1 − 3.4 can be characterized as two types: Type 1, in which there is a distinct and poorly conducting tissue element separating two layers; Type 2, in which there is no distinctive tissue element separating adjacent layers. The cornea epithelium and RPE provide examples of Type 1 boundary layers. Type 1 boundaries can be thought of as playing a role similar to that of cell membranes in electrical models of cells, though cell membrane conductances are usually much lower. Type 2 boundaries are typically anatomically distinct structures or tissues whose conductivities are similar.

#### Cornea

For Shells 1 and 2, the conductance for current flowing from the aqueous humor into the cornea (*G*_12_) and from the cornea into the gel/tear layer (*G*_23_) are required. The overall conductance of the cornea for some species (but not mouse) have been measured by imposing potentials across isolated pieces of cornea of known area, and measuring the translayer current, or vice-versa, imposing currents and measuring the resultant potential drop (Table 1). Suppose such a measurement of an isolated cornea yielded the conductance estimate *G*_cornea_. It would be incorrect simply to apply the relation *G* = *g /*Δ*L* in this case, as the cornea is inhomogeneous in thickness (Figs. 3A, 4A inset), with the epithelium in particular having much less extracellular water space, and thus necessarily being less conductive than the stroma. The relation *G* = *g /*Δ*L* nonetheless provides useful bounds on the conductivity of the sublayers. Thus, assuming that the thin corneal endothelium is responsible for the conductance *G*_12_, the epithelium for *G*_23_ and the stroma primarily responsible for the conductivity *g*_2_, one has *g*_2_< *G*_cornea_/ Δ *L* and *G*_12_, *G*_23_ > *G*_cornea_. To take advantage of these constraints, we took the approach of assigning different fractions of the total corneal resistance to each of its sublayers (Table 1).

#### Retinal pigment epithelium

At the posterior pole of the eye is the highly resistant RPE layer (with thickness ∼ 7 μm in mouse), whose tight junctions are a major component of the posterior blood-retinal barrier. The RPE is well characterized as a Type 1 boundary conductance: thus, for *θ* ≥ *θ*_3_, *G*_12_ (*θ*) = 0.005 S cm^-2^ (Fernandez-Godino et al., 2016).

As an example of a Type 2 boundary, consider the conductance *G*_23_ governing source/sink current between Shells 2 and 3 (Eq 3.3). Given the conductivity of Shell 2 as *g*_2_ (*θ*) and the distance from the center of Shell 2 to *r*_3_, the inner boundary of Shell 3, as *L* = Δ *r*_2_ / 2, the boundary layer conductance is estimated as *G*_23_ (*θ*) = *g*_2_ (*θ*) / (Δ *r*_2_ / 2) .

#### Converting boundary source/sink currents to volume current densities

Eqs. 3.1 − 3.4 are to be solved to determine the potential *V*. The units of the LHS of these equations is that of a volume current density (A cm^-3^), and the source/sink terms must also be expressed in the same units. To see how this is done, consider that the density of the current (A cm^-2^) flowing radially between Shells 1 and 2 at the polar angle *θ* is expressible as

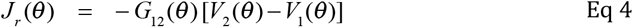

This current flows across a surface element of area Δ *S* at radial distance *r*_2_, the outer boundary of Shell 1 and inner boundary of Shell 2. In Eq 3.1, this current is leaving an element of volume Δ *vol*_1_(*θ*) = Δ *S*(*θ*) Δ *r*_1_, and so the total current leaving through the surface element is − *J*_*r*_ (*θ*) Δ *S*(*θ*) (units: A). Dividing the latter by the volume element gives the RHS of Eq 3.1. The volume element in Shell 2 into which the same current flows is Δ *vol*(*θ*) = Δ *S*(*θ*) Δ *r*_2_, and so the corresponding term in Eq 3.2 has Δ *r*_2_ in the denominator. The key idea is that the source/sink terms must be constructed so as to insure that the total current moving across any boundary element is conserved, i.e. satisfies the Equation of Continuity (Eq 2).

### Boundary conditions

Equations 3.1 – 3.4 are second order in *θ* and solving them therefore requires two boundary conditions (BCs) in *θ* for each equation. These BC’s express the absence of flux at the north and south poles:

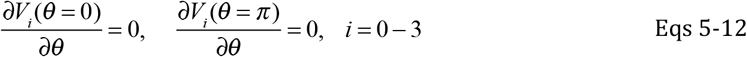

Equation 3.4 is also second order in *r* and thus requires two BCs in *r*, one each at the inner and outer radial boundaries of Shell 4. At the inner boundary (*r = r*_3_) we have

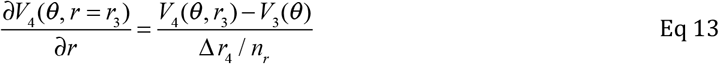

with Δ *r*_4_ = *r*_4_ − *r*_3_, while at the outer boundary we assume a no-flux condition:

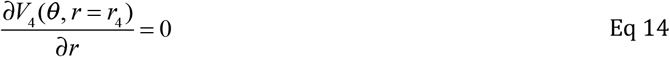

The former BC (Eq 13) is incorporated into the source term of Eq 3.4. The latter (Eq 14) is not strictly physically defensible, inasmuch as current can in principle flow beyond *r*_4_ in the cranium. The assumption can be rationalized post hoc, however, by showing that the potential approaching *r*_4_ is negligibly small.

### Solution of the 4-Shell Model Equations by Method of False Transients

Solutions to Eqs 3.1 – 3.4 were obtained with the Method of False Transients, as developed by (Northrop et al., 2013). Applied to the general version of Eq 3, this approach generates a solution of the pseudo-time-dependent partial differential equation (PDE)

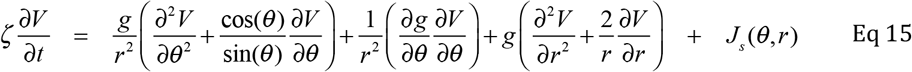

subject to the specified boundary conditions and a reasonable set of initial conditions. In Eq 15, *ζ* (units: S cm^-3^) is a scaling constant that serves to equate the dimensions on both sides of the equation, and which can be used to optimize the speed of convergence to zero. If well-behaved, the time-dependent solution evolves to a steady-state that is a solution of Eqs 3.1 – 3.4. The numerical method employed to integrate Eq 15 was the Numerical Method of Lines, a highly effective and stable method well described in the literature of partial differential equations (PDEs) (Schiesser, 1991; Schiesser, 2009). This method reformulates the PDEs into a series of ordinary differential equations (ODEs) by employing analytical approximations of the spatial derivatives at a series of discrete locations of the coordinates, and then numerically integrates the resultant ODEs with a standard integrator.

### Conservation of source/sink current

There is no formal guarantee that the Method of False Transients will yield a solution to the PDEs Eqs 3.1–3.4 that is unique and independent of the somewhat arbitrary initial conditions, and so experimentation with different plausible initial conditions was necessary. As the 4-Shell model is a closed system, we required that solutions satisfy global electrical conservation: thus, spatially integrated over the retinal surface, the current flowing outward from the photoreceptor or RPE source and the current returning to the corresponding sink was required to sum spatially to zero. For most conditions examined and reported here, convergence of Eq 15 to steady-state occurred within 50 or so iterative cycles, and conservation (zero net electrical flux) within each shell was satisfied to within 0.2%. Except for Shell 4, conservation can be assessed by plotting the current flowing in the shell as a function of the angular coordinate: this current: this plot must asymptote to zero at the south pole, *θ* = 180 deg. Electrical conservation in the 4-Shell Model is equivalent to the familiar constraint that must be obeyed by all (including spatially extended) models of electrically active cells, and follows from the Equation of Continuity, Eq. 2.

### Polar angle-dependence of a retinal layer source and layer resistances

The layers of the retina vary in thickness as a function of eccentricity, i.e., with distance from the optic nerve head. A classic example is the thickness of the outer nuclear layer (ONL; Fig. 4C), universally used to quantify rod photoreceptors in normal and diseased states. The local density of rods can be reasoned to be directly proportional to the ONL thickness, given that the rod cell body size is approximately constant: thus, in a small volume of retina, the total number of rods contained is equal to the ONL thickness multiplied by the surface area of the volume, divided by the effective volume of the cell body (the cell body multiplied by the packing fraction). Given that the rod density is proportional to the ONL thickness, the latter can be used as a surrogate for the source current density of the rod dark current as a function of eccentricity (compare Figs. 4C and 4D).

### Polar dependence of conductivity and conductance

The polar angle-dependence of the conductivities *g*_1_(*θ*) and *g*_2_ (*θ*) of Shells 2 and 3 necessitate inclusion of terms in Eqs 3.2 and 3.3 with the derivatives of the conductivities with respect to *θ* . The model as presented in Fig. 4 implicitly includes step changes in conductivity at each of a series of polar angles boundaries. Since step changes cannot be accommodated by the numerical integration methods we employed, we developed analytical expressions for the polar variation in conductivity based on logistic functions. The function describing a steplike change from conductivity level *g*_*n*_ to level *g*_*n*+1_ centered at angle *θ*_0_ is expressed as

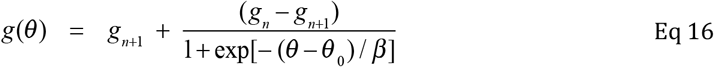

The parameter β determines the steepness of the transition and was set to 2– 3 degrees. Equation 16 can be readily generalized to accommodate a series of such changes (Fig. 4). This class of functions makes it possible to efficiently encode a polar distribution of conductivity with steplike changes that are analytically differentiable, as required by Eqs. 3.2 and 3.3.

## RESULTS

### Intraocular pressure varies about 25% over an hour-long experiment and ERG stability is maintained

The ERG affords the possibility of non-invasively assessing physiological function of the neural and electrically responsive glial and epithelial tissues of the mouse eye over an extended period of time. A potential problem is that anesthesia or mydriatic drugs used to dilate the pupil could cause changes in intraocular pressure (Kim et al., 2012), altering neural, glial or epithelial electrical function. The effect of the mydriatic cocktail used in the ERG experiments on intraocular pressure (IOP) was assessed by measuring IOP every fifteen minutes for an hour, with measurements also in a control group whose eyes were treated only with PBS (Fig. 5A). In parallel experiments the stability of the ERG was assessed by recording the response to a saturating flash every 5 min for 70 min (Fig. 5C, D), extracting the amplitudes the *a*-wave and the *c*-wave from the traces and plotting them as function of the time in the session (Fig. 5B). After a slight initial drop, IOP increased gradually over the hour to 23% above the average baseline value of 15.5 mmHg. The maximum value (19 ± 1.5 mm Hg), though statistically above the baseline, was not reliably different from the maximum (18.2 ± 1.5 mM Hg) of normal diurnal variation in one previous study (Aihara et al., 2003). It was also within error of the upper limit (∼ 17 mm Hg) of the range of IOP measured over 3 weeks in saline-injected control mice in another study (Sappington *et al*., 2010) that employed the same IOP measurement methodology as used in this study.

**Figure 5.**
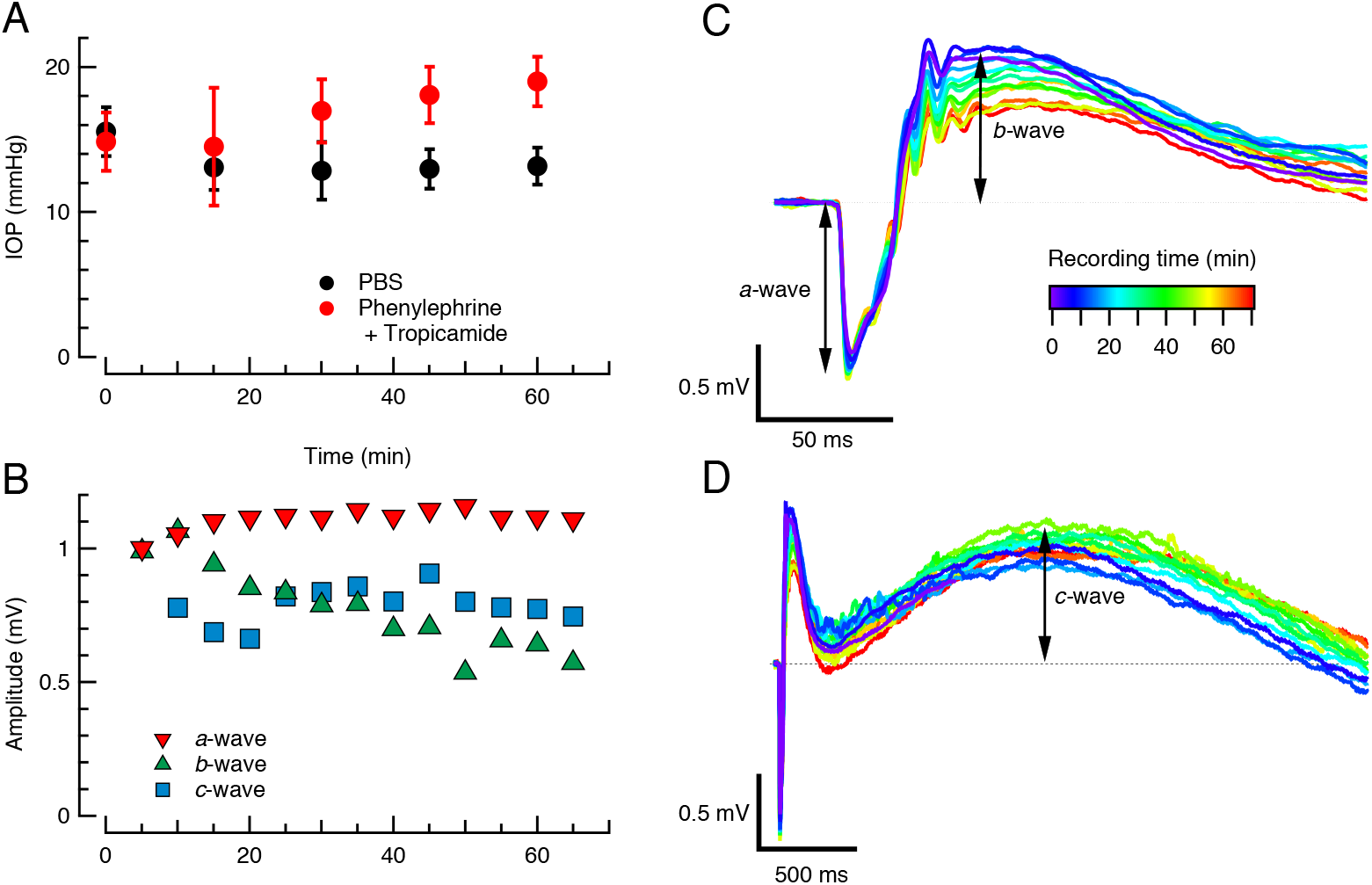
Stability of intraocular pressure and ERG recordings over a recording session. **A**. Intraocular pressure of two groups of anesthetized C57Bl6 mice in a 1-hr session: the eyes of one group (red symbols, mean ± SEM) were treated with the mydriatic protocol (Phenylalanine + Tropicamide) used in the ERG experiments, while the eyes of the control group received only PBS to maintain hydration. **B**. Amplitude of *a*-wave and *c*-wave components of the ERGs in response to a 500 nm saturating flash that delivered 2.6 ×10^5^ photons μm^-2^ to the retina as a function of time in the recording session. **C**. ERGs corresponding to the points in panel B; the traces are color-coded for the time at which they were measured in the session. **D**. Same set of ERGs as in C, but presented on much longer time base to show the *c*-waves.

The amplitude of the ERG *a*-wave was highly stable over the 70 min recording session after increasing 10% in the initial 15 min (Fig. 5C). The *c*-wave amplitude was less stable, declining by 30% in the initial 20 min, recovering to 85% to 90% of its baseline amplitude, and then maintaining ∼75% of its initial amplitude over the final 20 min of the session. We believe that the increase in *a*-wave amplitude arose from increased electrical isolation of the corneal apex (examined in the following section), but have no ready explanation of the *c*-wave change. Nonetheless for the initial decline, the *c*-wave was stable over most of the recording session, allowing reliable. measurement with stimuli of varied intensity. Overall, we conclude that the combined anesthetic, mydriatic and temperature-maintenance protocol affords good stability of ERG *a*-waves and *c*-waves over the hour-long recording epoch used in these studies.

### Conductive gel placed over the whole ocular surface greatly reduces the ERG amplitude

The saturating amplitude of the *a*-wave obtained in our standard recording condition (Fig. 1A-C) is routinely about 1 mV, 2- to 3-fold larger than in most recent mouse studies (Table S1). In the effort to pinpoint the basis of this disparity, we performed an experiment in which conducting gel was added to the surface of the eye external to the corneal ring electrode. This manipulation approximates the recording condition in ERG experiments in which the entire corneal surface is covered with a conducting medium. The addition of gel external to the ring electrode had a dramatic effect, reducing the ERG *a*-wave amplitude to 38% ± 0.02% (mean ± 95% CI, *n* = 8) of that recorded immediately prior to adding the extra gel, while producing minimal change in the ERG kinetics (Fig. 6A, B, E).

**Figure 6.**
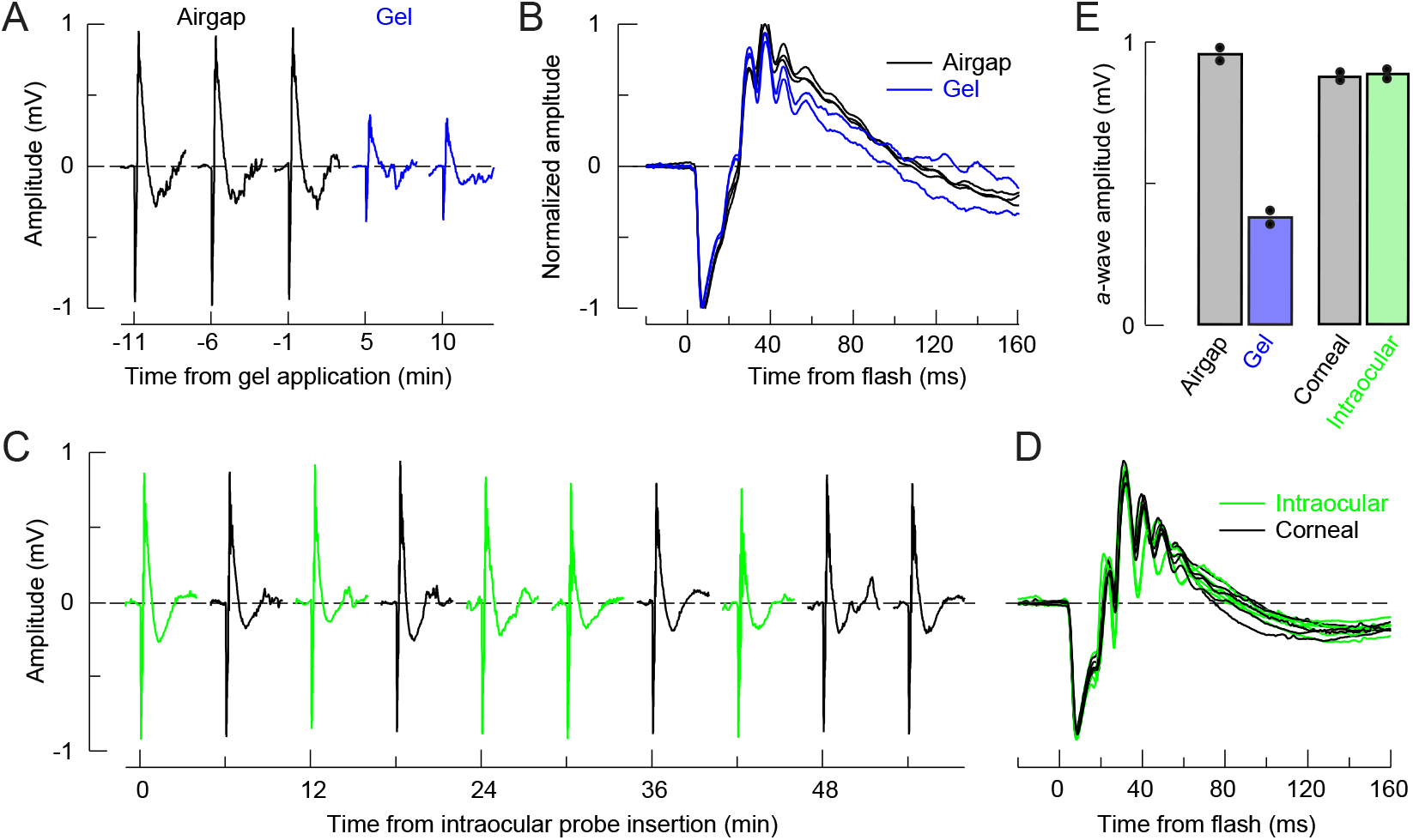
Manipulations of the ERG revealing inadequacies of the electric circuit model and the experimental basis for high amplitude recordings. **A**. Series of ERGs recorded under standard conditions to a saturating flash delivered 2.6 ×10^5^ photons μm^-2^ to the retina before (black traces) and after (blue traces) addition of conducting gel to the surface of the eye external to the corneal ring electrode. **B**. Traces from A superimposed and normalized to the maximal amplitude and presented on a brief time scale to illustrate the kinetics of the responses. **C**. Series of ERGs recorded in response to a saturating flash recorded with either a corneal electrode (black traces) or an intraocular electrode insert near the limbus (green traces; cf. Figs. 2A, 7B). **D**. Traces from C superimposed and presented on an expanded time scale. **E**. Summary bar chart: the points plot average from two replications of each experiment (A, C); each symbol plots the average of 4 or more responses in each condition, as in A, C. (cf. Tables 3, 4 for additional details.)

### The ERG has the same amplitude with a corneal and an intraocular electrode

A second telltale manipulation was performed by recording the ERG in the air-gap configuration with both the corneal electrode, and a second electrode that was inserted into the aqueous humor (Fig. 6C, D). Remarkably, the amplitude of the ERG recorded intraocularly and that recorded with the corneal electrode in the “air-gap” configuration were not materially different (Fig. 6C, D): specifically, the ratio of the saturating *a*-waves recorded with the corneal and intraocular electrodes was 0.98 ± 0.04 (mean ± 95% CI, *n* = 7; Fig. 6E; *p* > 0.25 for test of ratio against unity). This result implies a negligible drop in potential across the cornea in our standard recording condition, which creates an “air gap” external to the corneal ring electrode.

**Table 3:** Table 1. *a*-wave and *c*-wave amplitudes recorded in the “tears” and airgap configurations.

| | Mouse | “Tears” ($\mu\text{V}$) | Air gap ($\mu\text{V}$) | Ratio |
| --- | --- | --- | --- | --- |
| *a*-wave | 8362 | 390 | 965 | 0.40 |
|  | 8363 | 335 | 893 | 0.38 |
|  | 8364 | 290 | 877 | 0.33 |
|  | 7870 | 345 | 945 | 0.37 |
|  | 16076 | 338 | 945 | 0.36 |
|  | 16077 | 334 | 820 | 0.41 |
|  | 16078 | 343 | 850 | 0.40 |
|  | 16079 | 350 | 940 | 0.37 |
|  | **Average** | **341** | **904** | **0.38** |
|  | **SEM** | **10** | **20** | **0.01** |
| *c*-wave | 16076 | 127 | 660 | 0.19 |
|  | 16077 | 155 | 600 | 0.26 |
|  | 16078 | 220 | 713 | 0.31 |
|  | 16079 | 220 | 845 | 0.26 |
|  | **Average** | **181** | **705** | **0.25** |
|  | **SEM** | **27** | **60** | **0.03** |

**Table 4:** *a*-wave amplitudes recorded at the cornea or intraocularly in the air-gap configuration.

| Mouse | Corneal ($\mu\text{V}$) | Intraocular ($\mu\text{V}$) | Ratio |
| --- | --- | --- | --- |
| 8573 | 898 | 898 | 1.00 |
|  | 888 | 870 | 1.02 |
|  | 895 | 894 | 1.00 |
|  | 888 | 869 | 1.02 |
| 8570 | 897 | 982 | 0.91 |
|  | 898 | 940 | 0.96 |
|  | 874 | 914 | 0.96 |
| **Average** | **891** | **910** | **0.98** |
| **SEM** | **4** | **16** | **0.02** |

### Amplitude Reduction by Gel Is Inconsistent with the Circuit Model of the ERG

The Circuit Model (Fig. 3B) predicts the ratio of the corneally recorded *a*-wave and *c*-wave amplitudes relative to their respective source potentials as a ratio of resistances in the circuit. Thus, identifying *V*_PRL_ as the potential across the photoreceptor layer generated by the photoreceptor dark current, and *a*_max,airgap_ as the amplitude of the *a*-wave, we have

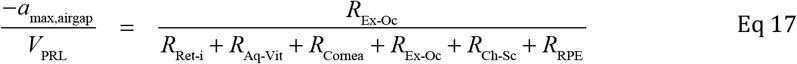

*V*_PRL_ is the difference between the potential at the photoreceptor synaptic layer (the outer plexiform layer, OPL) and that at the outer segment tips, and is positive, as the dark current flows out the inner segment and into the outer (Hagins et al., 1970). Taken as the suppression of the rod dark current, the *a*-wave has the opposite sign. The resistance *R*_Ex-Oc_ is that between the corneal electrode and the reference electrode. Similarly, for a transepithelial layer RPE source with potential *V*_RPE_ the Circuit Model predicts

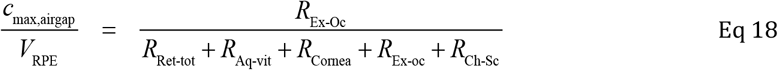

A problem with testing the predictions of Eq 17-18 against ERG data is that several of the resistances in these equations are difficult to specify, in particular the critical resistance *R*_Ex-Oc_, and moreover source amplitudes (*V*_PRL_, *V*_RPE_) must be specified. Nonetheless, a “source-independent” prediction can made by the circuit model for the effect of application of gel. When conducting gel is applied to the surface of the eye external to the corneal ring electrode, it is predicted to reduce the extraocular resistance to a lower value, *α R*_Ex-Oc_, with 0 < *α* < 1, and the reduction of the *a*-wave and *c*-wave amplitudes by the gel should be essentially the same (Appendix, Eq. A8). This prediction fails, as the *a*-wave is reduced by the factor 0.377 ± 0.023 (mean ± 95% confidence interval), while for the *c*-wave the reduction factor measured in the same experiments is 0.255 ± 0.076, a reliably different ratio (*p* < 0.0002). Additional predictions of the ERG Circuit Model can be tested provided specific values (or well-defined ranges of values) are assigned to the resistances. Subsequently, we present such tests, but find it helpful to that effort to present first the predictions of the 4-Shell Model, and then compare the two models’ predictions of the same data.

### The 4-Shell Volume Current Model Quantitatively Predicts the Effect of Air-gap vs Gel Conditions

A key feature of a volume current framework is that it allows current driven by a vitreal-positive potential to exit the intraocular media through several spatially distributed, parallel paths: these paths include that across the cornea described in the Circuit Model (Fig. 3B), but also a path laterally through the corneal stroma, and a third path through the tissues of the limbus and ora serrata (Fig. 4B). Qualitatively, a volume current framework can explain the effect of gel addition external to the corneal ring electrode as follows. In our standard recording condition the air gap external to the ring corneal electrode forces current traversing the corneal endothelium to exit through the weakly conducting corneal stroma (Table 1), enabling the potential at the corneal apex to remain relatively high, with current return to the PRL sink dominated by paths through the limbus and posterior extraocular fluid and tissue. In contrast, in the gel condition, the entire corneal surface becomes a massive parallel shunt path to the conjunctiva and posterior eye, “dragging” the corneal potential down.

These qualitative ideas are borne out quantitatively in solutions to the equations governing the potential distribution in the 4-Shell model for both PRL (Fig. 7) and RPE (Fig. 8) sources. For the standard recording condition with an airgap external to the corneal ring electrode the potential at the cornea apex internal to the ring is predicted to be essentially the same as that measured with an intra-ocular electrode (Fig. 7A, B). When conducting gel is applied to the whole ocular surface, however, the corneal potential is “pulled down” to the potential of the extraocular medium posterior to the eye (Fig. 7C, D), whose high conductivity insures its potential is relatively close to that of the PRL sink at the photoreceptors outer segments. The ratio of the *a*-wave amplitudes predicted by the 4-Shell Model for the two experimental conditions nearly perfectly matches that of the observations (Fig. 9A). Similarly, the 4-Shell Model predicts the effects of gel application to the eye in reducing the *c-*wave amplitude (Fig. 8), and moreover accounts for the relatively greater reduction of the *c-*wave vs. the *a-*wave (Fig. 9A). Finally, the 4-Shell Model nicely accounts for the nearly unity ratio of the *a*-wave amplitude recorded with a corneal electrode to that recorded intraocularly (Fig. 6; Fig. 9A).

**Figure 7.**
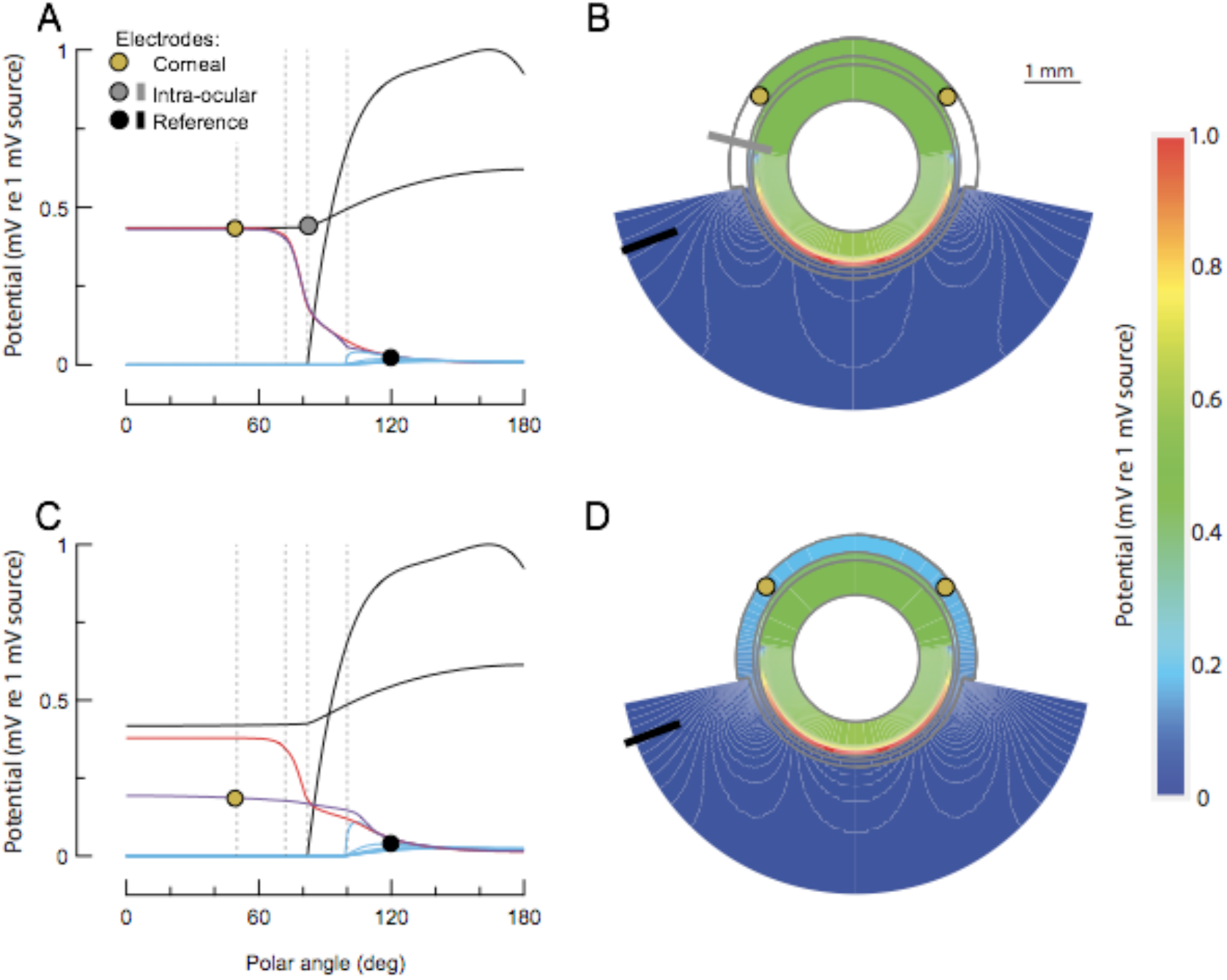
Predicted potential distributions in the eye and extraocular tissue for a 1 mV trans-photoreceptor layer source with (A, B), and without (C, D), an air gap on the cornea external to the ring electrode. **A**. Predictions for the air-gap recording condition, for recordings with the corneal electrode and an intra-ocular electrode. The traces are as follows: PRL source distribution (assumed to be 1 mV at its maximum near the optic nerve head), black curve peaking at 1 mV; intraocular media (Shell 1), thin black trace; Shell 2, purple trace; Shell 3, red trace; Shell 4, cyan traces (spaced radially at 0.44 mm). The predicted amplitudes are given by the differences between the corneal ring electrode (gold dot) or intraocular electrod (gray dot) and the reference electrode (black dot): *V*_corneal_ – *V*_ref_ = 0.386 mV; *V*_intraocular_ – *V*_ref_ = 40 mV. **B**. Colorimetric map of the potentials in A over the 4 shells (vertical gray lines define the boundary between layers in the anterior region, and illustrate isopotential contours (?) in the 4^th^ shell (extraocular tissue). **C**. Prediction for recordings made with conducting gel on the cornea outside the corneal electrode; *V*_corneal_ – *V*_ref_ = 0.11 mV. **D**. Topographic colorimetric map of the potential distribution in C (cf Fig. 4A). (Intraocular measurements could not be made when the whole cornea was covered with conductive gel.)

**Figure 8.**
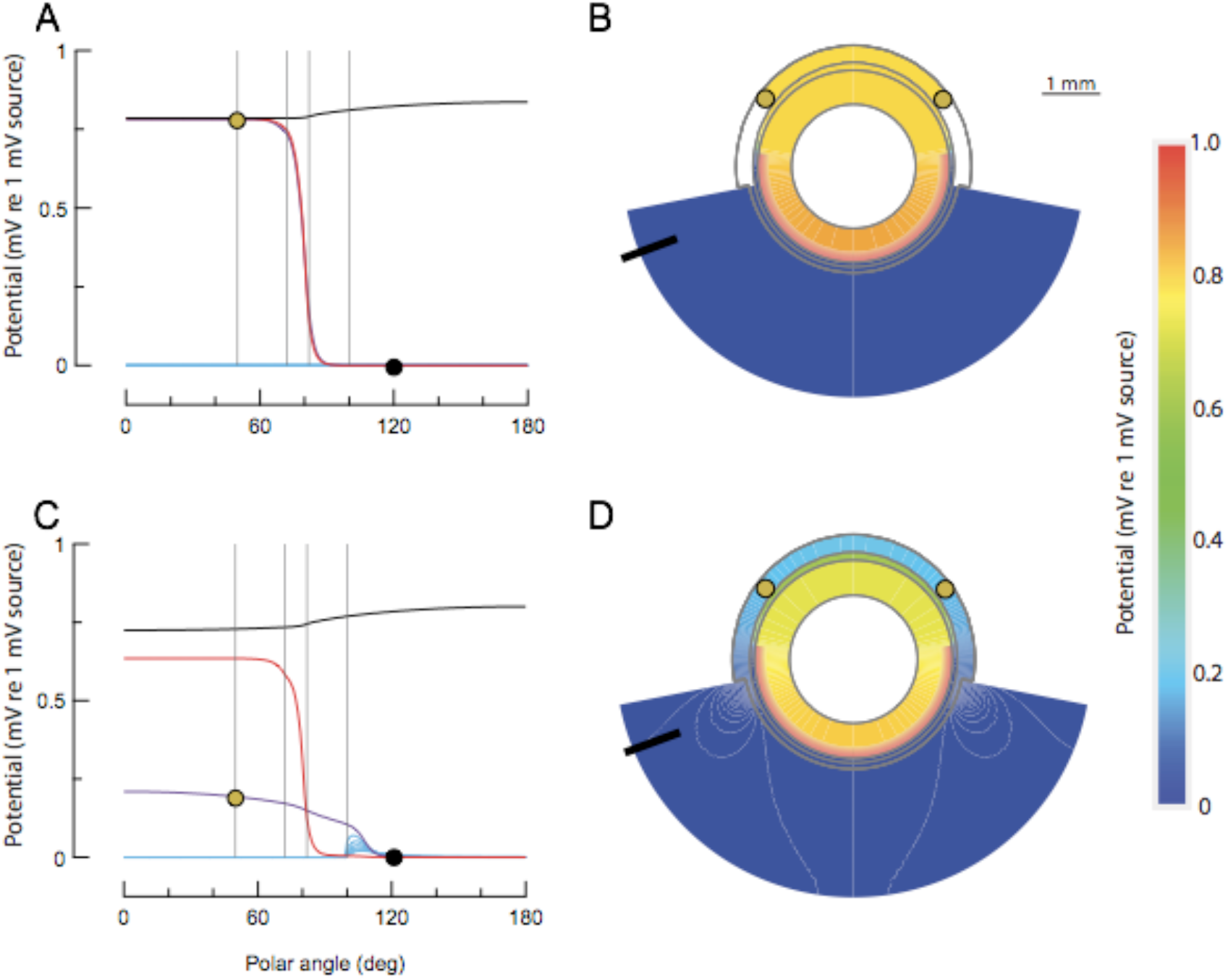
Predicted potential distributions for the RPE layer source with (A, B), and without (C, D), an air gap on the cornea external to the ring electrode. **A**. Predictions for the air-gap recording condition, for recordings with the corneal electrode and an intra-ocular electrode: the predicted amplitudes are the differences between the anterior electrode (gold dot, corneal; gray dot, intraocular) and the reference electrode (black dot): *V*_cornea_ – *V*_ref_ = 0.78 mV; *V*_intraocular_ – *V*_ref_ = 0.8 mV. **B**. Colorimetric map of the potentials in A over the 4 shells (the vertical gray lines define the boundary between layers in the anterior region. **C**. Prediction for recordings made with conducting gel on the cornea outside the corneal electrode; V_corneal_ – V_ref_ = 0.22 mV. **D**. Topographic colorimetric map of the potential distribution in C.

**Figure 9.**
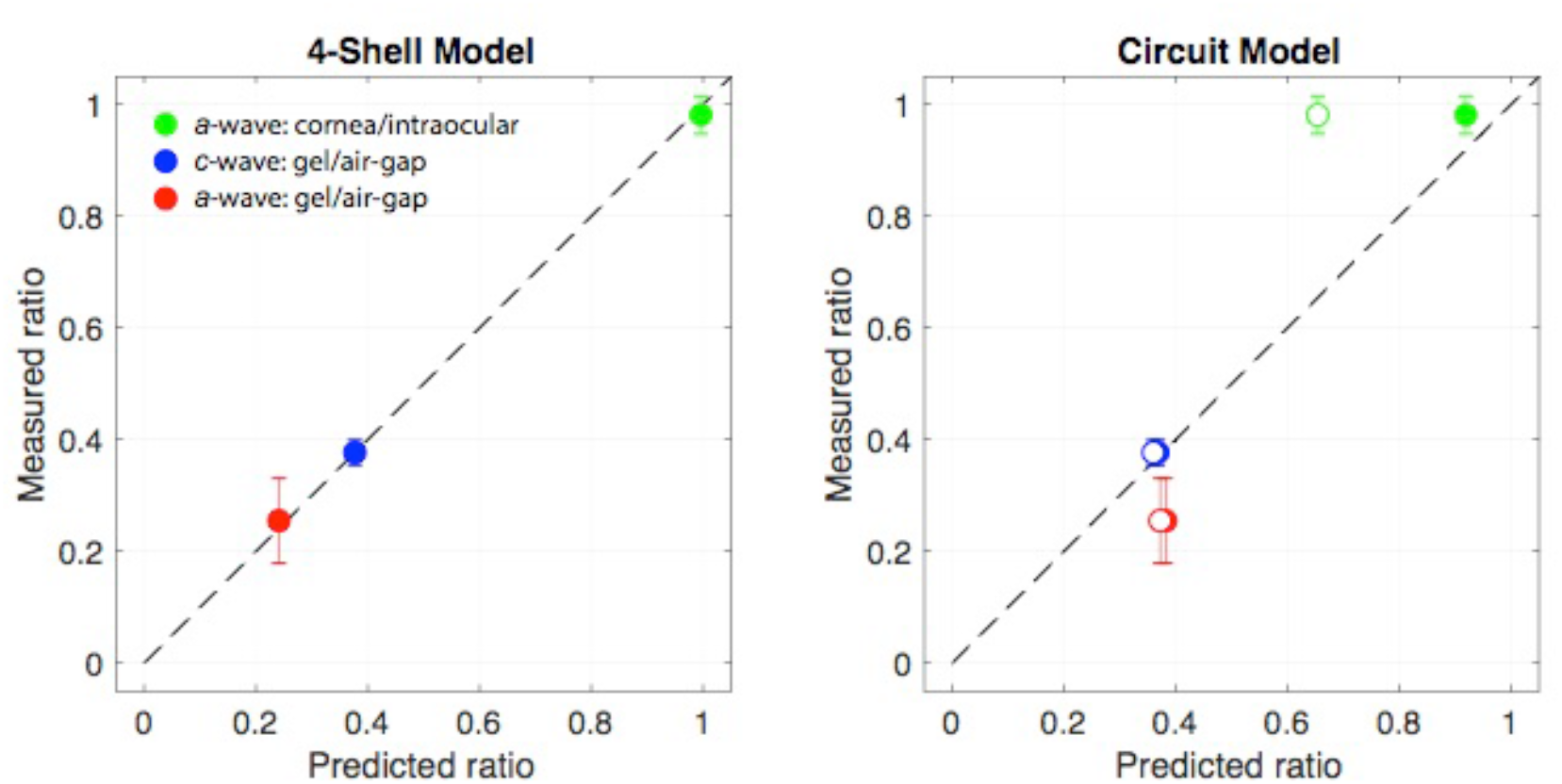
Comparison of measured potential ratios (Eq 19) with maximum likelihood predictions of the two models for different ERG components and recording conditions. **A**. 4-Shell Volume Current Model. Nearly perfect predictions, as indicated by the points falling on the unity-sloped line (dashed); error bars are 95% confidence intervals. **B**. Circuit Model (cf. Fig. 3B). The circuit resistances (Fig. 4B; Table 2) used to predict the ratios are maximum likelihood estimates obtained for two cases: (i) layer resistances restricted to be close to empirical measurements, and medium resistances restricted to be lower than rational upper bounds (open symbols); (ii) resistances with essentially unrestricted ranges. In both cases the predictions of the three ratios (Eq 19) fail with high statistical reliability (see text).

### The Circuit Model Fails to Predict Three Measured Source-Independent Ratios Correctly Predicted by the 4-Shell Model

To test the Circuit Model specific values needed to be assigned to its resistances. A perennial problem with the model has been its lack of specification of the extra-ocular resistance over which the ERGs are hypothetically measured (Fig. 4B). Data on the extra-ocular fluid and tissue conductivities (Table 1) provided a biophysical basis for assigning values or at least rationalizing ranges for the circuit’s resistances (Table S2). To do this we generalized the extra-ocular resistance (Fig. 3B) to include a serial element for the superficial medium on the cornea followed by two parallel resistive paths comprising the post-ocular orbital fluid and the post-ocular tissue, respectively. We then applied a search procedure (Appendix) to find maximum likelihood estimates (MLEs) of the resistances of the Circuit Model for predicting the three measured dimensionless ratios:

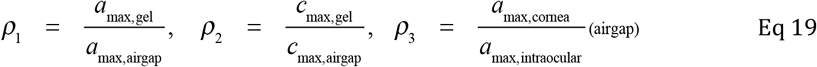

Because the source terms are the same for the numerator and denominator underlying each of the ratios *ρ*, *i* = 1− 3, the predictions for Eq 19 are source amplitude-independent (cf. Appendix). MLE predictions of the Circuit Model are provided for two cases (Fig. 9B): (i) essentially unrestricted ranges for all parameters; (ii) the ranges of resistances comprising the extra-ocular components restricted by rational upper bounds, and the ranges of resistances for which there are “tight” measurements restricted to ± 20% of the measurement. Both versions fail (Fig. 9B).

The maximum likelihood method provides a gauge for the comparing the Circuit and the 4-Shell models’ ability to predict the three source-independent ratios (Eq 19). Specifically, the parameter values that maximize the likelihood are those that minimize the weighted sum of squared errors, which is in effect a *χ* ^2^ statistic (Eq A15). For the Circuit Model with MLEs obtained from “restricted ranges” *χ* ^2^ = 588, while for the “unrestricted range” *χ* ^2^ = 77.1. The lowest possible value of the *χ* ^2^ statistic is 19, obtained when the predicted values of *ρ*_*i*_ exactly match the mean values of the ratios from the 19 empirical measurements. Thus, the Circuit Model fails quantitatively, even when its resistances are chosen as the optimal values from very unrealistic ranges. In contrast to these quantitative failures of the Circuit Model, the 4-Shell Model nearly perfectly predicts the three ratios, as demonstrated by the points falling very close to the slope unity line (Fig. 9A) and by the low value of the error statistic, *χ* ^2^ = 20.0.

## DISCUSSION

### Limitations and failures of the Rodieck-Ford Circuit of the Electroretinogram

The familiar Electrical Circuit Model of the ERG (Fig. 3B) was proposed by Rodieck and Ford (1969). The model was elaborated in Rodieck (1973, Fig. XVIII-2) and (Steinberg, 1985b), and has been widely used as an instructional device in basic and ophthalmic science -- e.g., (Perlman, 1995). In this investigation we found that the Circuit Model cannot account for several measureable, “source amplitude-independent” features of the mouse ERG (Fig 9B).

The Circuit Model also has notable conceptual limitations, some of which have been discussed before (Steinberg, 1985a). Perhaps the most obvious one is that it does provide a way to measure or calculate the extra-ocular resistance over which the ERG is measured (*R*_Ex-Oc_ in Fig. 3B): thus, the return current path for retinal and RPE sources has never been measured or specified in a manner that could locate a reference electrode in this path and allow the potential difference between it and the corneal electrode to be calculated. Here, we used analysis of the tissue conductivities and layer resistances (Table 1) to determine an effective resistance of the extraocular path, and found that even with maximum likelihood estimates of all the resistances, the Circuit Model failed (Fig. 9). The core problem is that the eye and extraocular tissue have a complex 3-dimensional structure, and understanding the return path for the sources of the ERG, and ultimately the amplitude of components of the ERG, requires analysis of the volume current flow in this structure.

### Successes of a 4-Shell Volume Current Model of the ERG

Applying the physical principles of tissue current source density (CSD) analysis (Eqs 1-2), we undertook to create a volume current model of the mouse ERG. Considerations of the shape of the eye and its azimuthal symmetry led us to formulate the model in spherical coordinates (Eq 3), and considerations of the conductivities and conductances of various component ocular and extraocular components (Table 1) suggested that a good approximation of the structure could be made with 4 concentric conducting shells surrounding a spherical lens (Fig. 4). The 4-Shell Model rests on physically measured radial and polar angle-dependent transitions of the eye, and on independent specification of the conductances and conductivities (Fig. 4E, F; Table 2). The 4-Shell Model allows the location of the reference electrode in the relevant volume to be specified (Figs. 2, 4B), as well as specification of polar angle-dependent distributions of different retinal sources (Fig. 4D), and enables systematic exploration of the conductivities of the ocular tissues and media.

The 4-Shell Model was able to provide a nearly perfect account of the “source-independent” ratiometric observations that the Circuit Model fails to explain (Fig. 9). Equally important as this quantitative success, the 4-Shell Model explains why conducting gel added to the eye greatly reduces the ERG amplitude, why this manipulation reduces the saturating *c*-wave amplitude more than that of the *a*-wave, and why the potential drop across the cornea is effectively negligible in the air-gap recording condition. Specifically, the 4-Shell Model reveals that in the “airgap” recording condition the relatively high potential at the corneal apex created by a PRL source results from negligible current flows across the corneal surface, so that in this condition current must flow within the cornea toward the sinks in the posterior eye through the relatively resistant corneal stroma (Fig. 7A, B). Addition of conducting gel to the entire ocular surface creates a low resistance shunt that allows current to exit the eye through the whole corneal surface, and to return more readily to the cellular sinks: with gel over the whole corneal surface the potential drop across the corneal apex is 2 to 3-fold lower, “pulled down” by the shunt’s electrical proximity to ground (Fig 7C, D). The explanation for the greater reduction of the *c*-wave than the *a*-wave by the application of gel has two components. First, the potential predicted to be measured at the cornea per mV RPE source in the airgap configuration is 2-fold greater than for a PRL source (compare Figs 7A and 8A). Second, for the RPE source, the reference electrode is effectively at the potential adjacent the sink, the RPE baso-lateral surface potential, while the PRL sink lies across the RPE layer. Finally, the 4-Shell Model explains why the intraocular and corneal electrodes measure the same potentials in the airgap configuration: in the absence of material current flow across the cornea, there can be no potential drop, as the two are inextricably linked (Eq. 2).

### Achieving the highest possible amplitude ERGs in mouse and variation in published amplitudes

The 4-Shell Model helps to explain why the airgap configuration produces ERGs with the highest amplitude in the literature, and in doing also so helps to explain the wide variation in amplitude in the literature (Table S1). As explained in the previous paragraph, the airgap outside the corneal ring electrode forces the potential at the corneal apex to match that of the underlying aqueous humor (Fig. 6). Common ERG experimental practice, however, floods the exposed extraocular surface with conductive medium, lowering the magnitude of potentials at the cornea, as shown here with gel addition, and explained by the 4-Shell Model (Figs. 7, 8).

Another practical insight provided by the 4-Shell Model is that the location of the reference electrode in extraocular tissue is largely irrelevant to the measured amplitude (Figs. 7-8). The reference electrode in mouse ERG recordings has been variously positioned subcutaneous to the dorsal scalp (as in our experiments), in the mouth (Lyubarsky and Pugh, 1996; Lyubarsky et al., 2004) and in other extraocular tissues. The 4-Shell Model reveals that the potentials generated by retinal sources in the extraocular tissues are negligibly above ground (Figs. 7, 8), and so optimum positioning of the reference electrode is principally a matter of stability, and minimizing electrical noise that may arise from extraocular (and other) muscles, and from breathing or heart-beat artifacts which can propagate through the body conductor.

### Challenges for the 4-Shell Model of Volume Conduction in the Eye and Extraocular Tissue

In this investigation a 4-Shell Model was tested against ratiometric “source amplitude-independent” predictions for PRL and RPE sources of unit (nominal 1 mV) amplitude. To achieve its full utility, the model should predict the absolute amplitude of the *a*-wave and *c*-wave based on a biophysical description of the sources. In two companion papers, we undertake to develop these descriptions, predict the absolute amplitude and kinetics of the source potentials based on the physiology of the underlying currents, and compare the predictions with measurements.

## Appendix

Evaluation of the Rodieck-Ford Electrical Circuit Model of the ERG

### Principles

The Rodieck-Ford circuit model of the ERG (Fig. 3B) is a closed loop electrical circuit. According to Kirchoff’s Voltage Law, the sum of the potential drops around the circuit from any starting point is zero. Defining *V*_PRL_ as the potential across the photoreceptor layer (PRL) arising from the rod dark current, it follows that the sum of the potential drops over all the other resistors in the circuit is –*V*_PRL_. We consider the two experimental manipulations of this study -- “airgap” and “gel-- and the potential between a corneal (or intraocular) electrode (Fig. 1) and the reference electrode whose location in the extra-ocular tissue is specified (Fig. 2). We treat the measured saturating *a*-wave corresponds to the potential change caused by the light-driven suppression of the rod dark current.

### Photoreceptor Layer Source

*Case 1*: *Airgap*. The potential across the PRL drives a current *I*_1_ around the circuit (with positive current flowing clockwise, and potentials across each resistor defined as the difference from the side more- to that less-clockwise). Then from Kirchoff’s Law

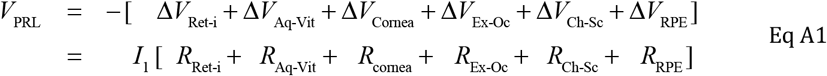

ERG recordings measure the difference in potential between an electrode on the extraocular face of the cornea and the reference electrode somewhere in the extra-ocular path. (The details of the extraocular path are murky in the model, but without loss of generality, assume that the reference electrode is located at the “south side” of *R*_Ex-Oc_.) Then, the saturating amplitude of the *a*-wave in the airgap case is

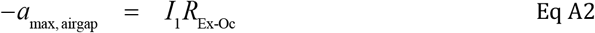

(The negative sign in front of *a* _max,airgap_ is required because the *a*-wave arises from the suppression of the PRL layer dark current that creates *V*_PRL_.) Combining Eqs 1 and 2, we have the ratiometric, dimensionless relation

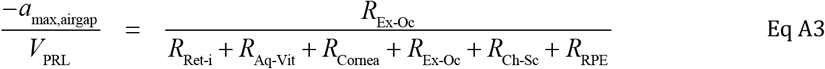

Note that the same current *I*_1_ is present in Eq A1 and Eq A2, so this term cancels out of Eq A3.

*Case 2: Gel*. The addition of gel to the surface of the eye outside the ring electrode reduces the resistance of the extra-ocular path to a value *αR*_Ex-Oc_, with 0 < *α* < 1 (again, the current, *I*_2_, different from *I*_1_, cancels out):

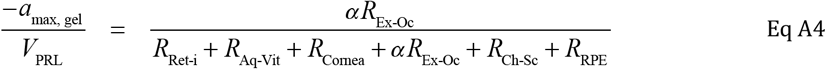

The measured ratio of the LHS of Eq A4 divided by the LHS of Eq A3 is 0.38 ± 0.01 (mean ± SEM). Thus, taking the ratio of Eqs A3 and A4 we have

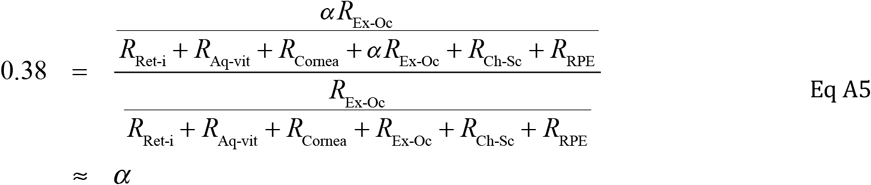

The second line of Eq A5 is rationalized on the grounds that the effective resistances of the cornea and RPE are dominant (and can be more closely associated with numerical values).

### RPE source

We can perform the same analysis as above for an RPE source, mutatis mutandis, i.e., keeping in mind that the source layer resistance is absent from the ratio of the total potential generated by the source around the circuit to the measured ERG component at the cornea. Thus, the equivalent of Eq A3 is

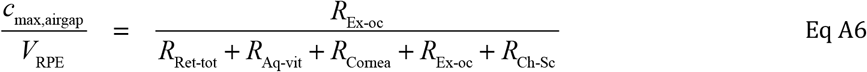

and the ratio for the gel condition is

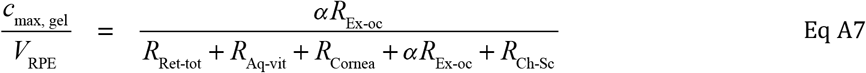

The measured reduction in c-wave amplitude with the addition of gel is 0.25 ± 0.03 (mean ± SEM). This observation then leads to the “double-double” ratio

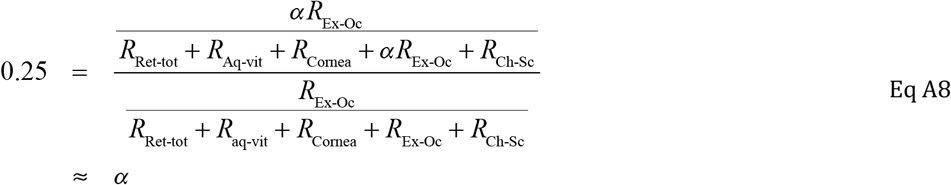

To obtain the second line of Eq A8 it is assumed that *R*_Ex-Oc_ ≪ *R*_Ret-tot_ + *R*_Aq-Vit_ + *R*_Cornea_ + *R*_Ch-Sc_ . This latter assumption is less readily defended than in the case of the PRL source, since the RPE is generally accepted as the major resistance in the circuit. However, consideration of estimates of most of the resistances in the circuit (Table S2) supports the assumption.

### Photoreceptor Layer Source: Corneal vs. Intraocular Electrode, Airgap Configuration

For an intraocular electrode, one has in parallel with Eq A2

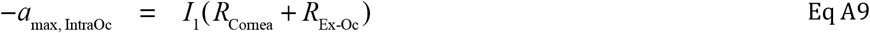

so that

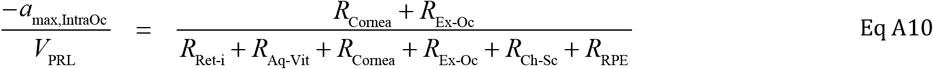

Dividing Eq A3 by Eq A10 then gives the predicted ratio of the *a*-wave recorded at the cornea vs. intraocularly:

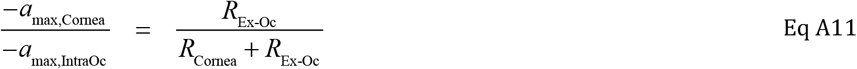

### Rejection of the Circuit Model

The measured fractional reductions of the amplitudes of the *a*-wave (0.377 ± 0.010, mean ± SEM, n = 12; Eq 5), and the *c*-wave (0.255 ± 0.028, n = 4; Eq 8) by the application of gel to the cornea outside the ring electrode are highly reliably different (*t*_df=10_ =5.78, *p* = 0.00018). Thus, fact that the fractional reduction *α* in the extracellular resistance by the addition of gel to the ocular surface exterior to the corneal ring electrode is different when estimated with ERGs from a PRL vs. RPE source is highly inconsistent with the circuit model. (The unequal n’s in the test above came about because in the initial experiments in which gel was added the duration of the recording epoch was too short to measure the c-wave. If only experiments in which both *a*-wave and *c*-wave were measured are used for the statistical test, the estimates of *α* were 0.385 ± 0.014 and 0.255 ± 0.028, *p* = 0.0028).

### Testing the assumptions

The assumptions made in Eq A5 and Eq A8 should be tested against available data. (In Companion Paper II, we estimate the resistances of most of the component resistors in the Rodieck-Ford circuit.)

### Maximum Likelihood Parameter Estimation and Hypothesis Testing

Define three empirical ratiometric statistics derived from ERG experiments:

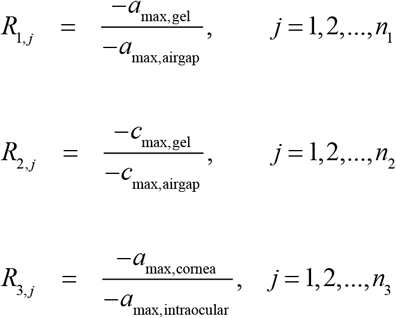

Here, the index identifies the *j*^th^ independent experiment from which the ratio is derived.

Let 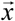 represent an n-dimensional vector of parameters for a parameter space for either of the models, RF Circuit or 4-Shell. Define the likelihood of the specific parameter 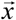 as follows:

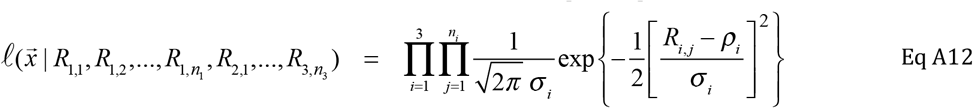

This assumes that the random variables *R*_*i, j*_ are distributed normally (i.e., with a Gaussian distribution) with mean *ρ*_*i*_ and standard deviation *σ* _*i*_, a generally acceptable assumption for any realistic random variable with a well defined mean and finite standard deviation. The product on the RHS of Eq A12 has a positive value less than unity, so taking the negative of its (natural) logarithm, gives rise to an all-positive function, such that when the original function is at its maximum, the negative logarithm will also be. Thus, the maximum likelihood estimate 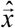 is the value in the parameter space 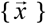 that maximizes the negative logarithm of the likelihood function. It is straightforward to show by differentiating the negative log likelihood function with respect to the parameters *ρ*_*i*_ and *σ* _*i*_, setting the results to zero and solving the equations that the maximum likelihood estimates from the given data set are

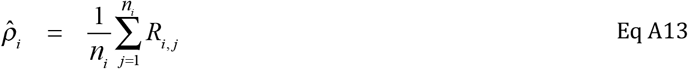

and

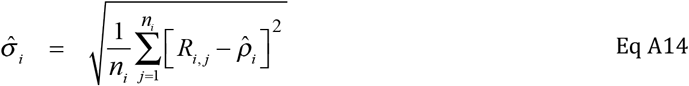

These results are not surprising: the MLEs are the sample mean and standard deviations. However, they also tell us how to compute an error function over the parameter space that allows the MLEs in that space to be determined. Specifically, consider the error function

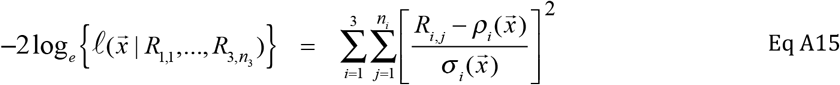

In other words, for each specific set of parameters, one first finds the MFT solution of the PDEs, and then computes the theoretical ratios 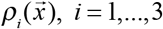. Next, one computes the values 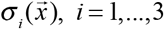 with Eq 13, and then Eq A15 gives the value of the error function at the parameter location 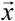. Note that the MLE vector 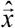 in the parameter space is precisely that which generates the prediction 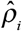 of Eq A13: thus, if one finds a parameter set 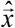 such that it produces 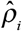 of Eq A13, then one has an MLE. Note also there is no guarantee that 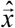 is unique: it could be a subspace of values, so one must be careful about any claims of uniqueness. However, if we scan, even crudely, a reasonable multidimensional rectangle and find that Eq A15 is always larger than the value that predicts the results in Fig. 9B, we can confidently assert that the value found is “the” MLE subject to the parameter space restrictions.

**Table A1:** Resistances for the Rodieck-Ford Circuit Model.

| *Layer/tissue/component* | Symbol | Thickness<br>($\mu\text{m}$) | Resistivity<br>($\Omega \text{ cm}$) | Resistance<br>($\Omega \text{ cm}^2$) | Circuit<br>Resistance<br>($\Omega$) |
| --- | --- | --- | --- | --- | --- |
| Retinal pigment epithelium (RPE) | $R_{\text{RPE}}$ | 6.0 | $3.3 \times 10^5$ | 200 | 1050 |
| Outer Retina (PRL) | $R_{\text{Ret-o}}$ | 105 | 5000 | 50 | 260 |
| Inner Retina | $R_{\text{Ret-i}}$ | 85 | 4760 | 40 | 210 |
| Aqueous & Vitreous | $R_{\text{Aq-Vit}}$ | 600 | 70 | Shell | 450 |
| Cornea | $R_{\text{Corn}}$ | 145 | $2.8 \times 10^4$ | 400 | 3175 |
| Combined ExtraOcular Path | $R_{\text{ExOc}}$ | -- | -- | -- | 1750 |
| Extraocular fluid, Cornea | $R_{\text{ExOcCorn}}$ | 100 | 60 | Shell | (750) |
| Extraocular fluid, Orbit | $R_{\text{ExOcOrb}}$ | 100 | 80 | Shell | (750) |
| Extraocular Tissue | $R_{\text{ExOcTiss}}$ | 400 | 80 | Shell | (225) |
| Choroid & Sclera | $R_{\text{Ch-Sc}}$ | 80 | 550 | 50 | 10 |
| Total | $R_{\text{Total}}$ | -- | -- | -- | 6905 |

*Table notes.* An entry in column 6 is obtained by multiplying that in column 5 of the same row by the tissue surface area, taken to be 19 mm<sup>2</sup> for the retina, and 12.6 mm<sup>2</sup> for the cornea, with the exception of entries identified as “Shell”. For these latter entries the component was treated as a hemispherical shell with radial thickness as given in column 3 (converted to cm), and resistance
calculated as $R = \int_{\theta_1}^{\theta_2} [\rho / A(z)] dz$ , where $\rho$ is resistivity (col 4), $z$ is an axial coordinate running
from the corneal apex to the ocular posterior pole, and $A(z)$ is the conductive cross-section at depth $z$ (see Supplement for Additional Details). In the posterior extra-ocular region, the conductance of the layer of extraocular fluid and that of the extraocular tissue were taken to be in parallel: thus, their individual resistances are given in parentheses, and only the parallel combination of the resistances was used in the circuit calculations. The superficial fluid layer bathing the cornea, however, was taken to be in series with the other extraocular resistances, and so the resistance of the “Combined ExtraOcular Path” is the sum of the resistance of the superficial fluid bathing the cornea, and the parallel combination of the resistances of the fluid in the orbit and of the extraocular tissue. In column 6 entries in parentheses are 50% of maximum possible resistances, as defined by application of the above formula.

Rodieck, R.W., and R.W. Ford. 1969. The cat local electroretinogram to incremental stimuli. *Vision Res*. 9:1-24.

## SUPPLEMENTARY MATERIAL

### Variability in ERG a-wave amplitudes amongst extant studies

Table S1 summarizes mouse ERG data from 10 well regarded studies of C57Bl/6 mice since investigations of mice employing corneal ERGs became commonplace. The average *a*-wave amplitude is about 50% that measured in the standard “air-gap” recording configuration of a previous (Peinado Allina et al., 2017) and the present investigation (last two rows of the table). As far as we can discern, all of the studies – with the possible exception of that by (Robson and Frishman, 2014) – flooded the cornea with conducting medium.

**Table S1:** Comparison of mouse corneal ERG *a*-wave amplitudes from different studies.

| Amplitude (μV) | Reference | Table/Figure in publication |
| --- | --- | --- |
| 464 | (Lyubarsky and Pugh, 1996) | Table 1 |
| 600 | (Kang Derwent et al., 2007) | Fig. 3 |
| 445 | (Hetling and Pepperberg, 1999) | Figs. 1, 6 (avg) |
| 350 | (Knop et al., 2008) | Fig. 4 |
| 500 | (Abd-El-Barr et al., 2009) | Fig. 2 |
| 450 | (Vinberg et al., 2014) | Fig. 2 |
| 800 | (Robson and Frishman, 2014) | Fig. 11 |
| 200 | (Shahi et al., 2017) | Fig. 4 |
| 590 | (Bush et al., 2019) | Fig. 1 |
| 350 | Kim et al. 2020 | Fig. 1 |
| 475 ± 156 | Mean ± SD |  |
| 1035 ± 55 (n=5) | Mean ± SD (Ronning et al., 2018) | Fig. 5 |
| 904 ± 56 (n=14) | Mean ± SD; this study | cf. Fig. 5, Table 3 |

*Table notes.* Amplitude of the saturating ERG *a*-wave reported in a collection of studies of C57Bl/6 mice from different laboratories over the past 25 years. The last two rows of the table summarize results from this laboratory, including the present.

---

**Fig. S1.**
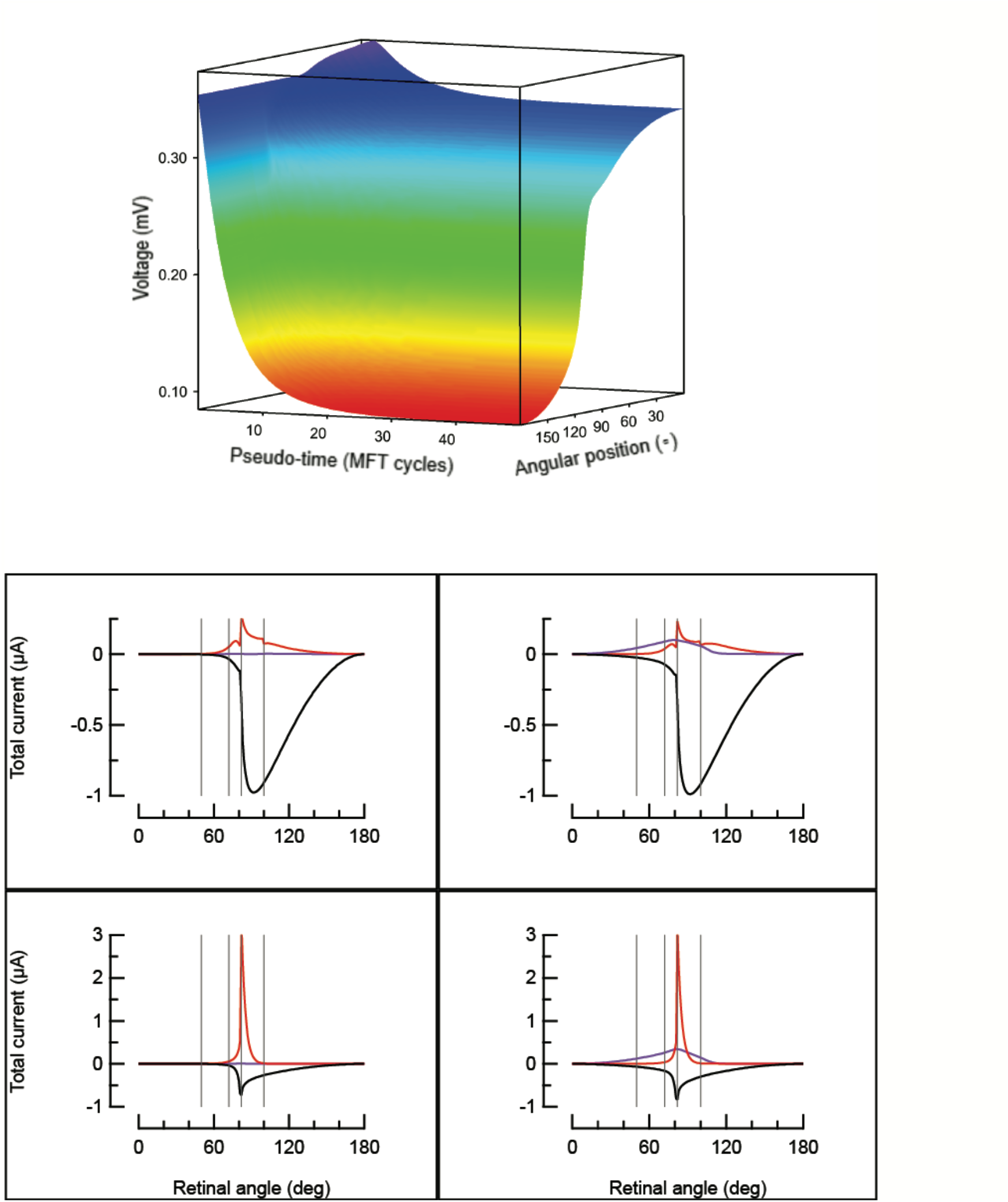
Solution of the Equation of Continuity (EOC) applied to the 4-shell model of the eye with the Method of False Transients (Eq. 15). Upper panel shows the potential distribution in the vitreous/aqueous shell (Fig. 4B, shell 1) as a function of the iteration cycle for a 1 mV trans-photoreceptor layer source, and demonstrates achievement of a highly stable result in 50 cycles with the final potential distribution the same as that in Fig. 7A. The lower 4 panels plot the current distributions in different shells on the angular coordinate, with the two panels in the first column representing results from the “airgap” configuration of the corneal electrode, and the two panels in the left column results from the “gel/tears” recording configuration. The large negative trace (black) is the current flowing in shell 1 (aqueous vitreous; the sign is negative because the current flows from the posterior pole toward the anterior pole of the eye); the red traces in the lower panels plot the current flowing in shell 2 (cornea/sclera), while the red traces in the upper panels the current in shell 3 (superficial fluid). Obedience to the EOC is demonstrated by the equal and opposite value of the total source and total sink currents, as was generally satisfied to within less than 0.01%.

